# Specification of human regional brain lineages using orthogonal gradients of WNT and SHH in organoids

**DOI:** 10.1101/2024.05.18.594828

**Authors:** Soraya Scuderi, Tae-Yun Kang, Alexandre Jourdon, Liang Yang, Feinan Wu, Alex Nelson, George M. Anderson, Jessica Mariani, Vivekananda Sarangi, Alexej Abyzov, Andre Levchenko, Flora M. Vaccarino

## Abstract

The repertory of neurons generated by progenitor cells depends on their location along antero-posterior and dorso-ventral axes of the neural tube. To understand if recreating those axes was sufficient to specify human brain neuronal diversity, we designed a mesofluidic device termed Duo-MAPS to expose induced pluripotent stem cells (iPSC) to concomitant orthogonal gradients of a posteriorizing and a ventralizing morphogen, activating WNT and SHH signaling, respectively. Comparison of single cell transcriptomes with fetal human brain revealed that Duo-MAPS-patterned organoids generated the major neuronal lineages of the forebrain, midbrain, and hindbrain. Morphogens crosstalk translated into early patterns of gene expression programs predicting the generation of specific brain lineages. Human iPSC lines from six different genetic backgrounds showed substantial differences in response to morphogens, suggesting that interindividual genomic and epigenomic variations could impact brain lineages formation. Morphogen gradients promise to be a key approach to model the brain in its entirety.

## Introduction

Decades of neurodevelopmental studies have demonstrated that the establishment of brain regions and the specification of cell fates are regulated by morphogen gradients released from different patterning centers along the neural tube ^1-3^. Morphogens, which include WNTs and sonic hedgehog (SHH), act in a concentration-dependent manner to trigger intracellular signaling pathways and activate specific transcription factors (TFs) along the axes of the neural tube ^4^. WNTs are posteriorizing morphogens since their release from the tail bud create a posterior-anterior gradient in the dorsal neuroectoderm, acting in complement to retinoic acid signaling ^5^. This gradient separates the forebrain (i.e. telencephalon and diencephalon), anteriorly, from the midbrain and the hindbrain (i.e., cerebellum, pons, medulla oblongata), posteriorly. Conversely, SHH is a ventralizing morphogen which diffuses from the notochord into the ventral neural tube to induce floor plate and ventral neuronal identities ^6,7^. In the telencephalon, this gradient distinguishes for instance the dorsal cortical plate from the ventral ganglionic eminences (GE), and in the midbrain, the ventral nuclei (such as the substantia nigra) from the dorsal optic tectum ^8^.

With advancements in stem cell technologies, a multitude of brain regions can be modeled *in vitro* by exposing embryonic or induced pluripotent stem cells (iPSCs) to specific concentrations of morphogens and other signaling molecules known to facilitate the specification a given region of the nervous system. In organoids, this has been successfully employed to generate optic vesicles, cortex, hippocampus, thalamus, or cerebellar fates in a 3D structure ^9-16^. However, only a few recently introduced methods sought to explore the effects of continuous morphogens gradients which can achieve intermediate concentrations of signaling molecules, as they occur in vivo ^17-19^. These methods have not yet explored the potentials of simultaneous orthogonal gradient fields along two axes, like the cooccurrence of posteriorizing and ventralizing gradients in vivo ^20^. Here, we developed a mesofluidic device that allowed us to expose seven control iPSC lines to two orthogonal morphogen gradients, mimicking simultaneously both the ventralizing action of SHH and the posteriorizing action of a WNT agonist. Using bulk and single-cell RNA sequencing (scRNA-seq) as well as protein-level characterization, we evaluated the dual gradients’ ability to recreate regional specification in organoids and control the formation of the diverse neuronal fates observed in the human brain. We find that the combination of WNT and SHH gradients generates stepwise patterns of transcriptional programs that specify a variety of cell types present in each major region of the brain. Furthermore, the responsiveness to these gradients differed across the seven iPSC lines, suggesting that early developmental patterning outcomes may depend on the genetic and epigenetic states of the donor cells.

## Results

### 1. Exposing organoids to a WNT/SHH orthogonal gradient reproduced early brain regional specification

To reproduce the diversity of human brain regions, we sought to expose human iPSC-derived embryoid bodies (EBs) to continuous gradients of morphogens during the earliest stage of neural induction. For this purpose, we developed Duo-MAPS, for Dual Orthogonal-Morphogen Assisted Patterning System, a mesofluidic device consisting of a closed culture area (5 communicating chambers delimited by octagonal pillars) flanked by sets of reservoirs (**Fig. 1A, Fig. S1A).** With Duo-MAPS as many as 200 EBs mixed in a collagen/agarose hydrogel can be embedded inside the culture area and be exposed simultaneously to two orthogonal morphogen gradients formed by passive diffusion. One morphogen is added to one of the large reservoirs creating a diffusion gradient along the horizontal axis of the device, while the other is added to a set of small side reservoirs creating a diffusion gradient along the vertical axis of the device. Following gradient exposure, EBs can be extracted separately from each chamber (numbered C1 to C5) to be further cultured and analyzed. The Duo-MAPS production and usage are easily scalable as it doesn’t require pumps or specific installation (**Methods**).

**Figure 1.**
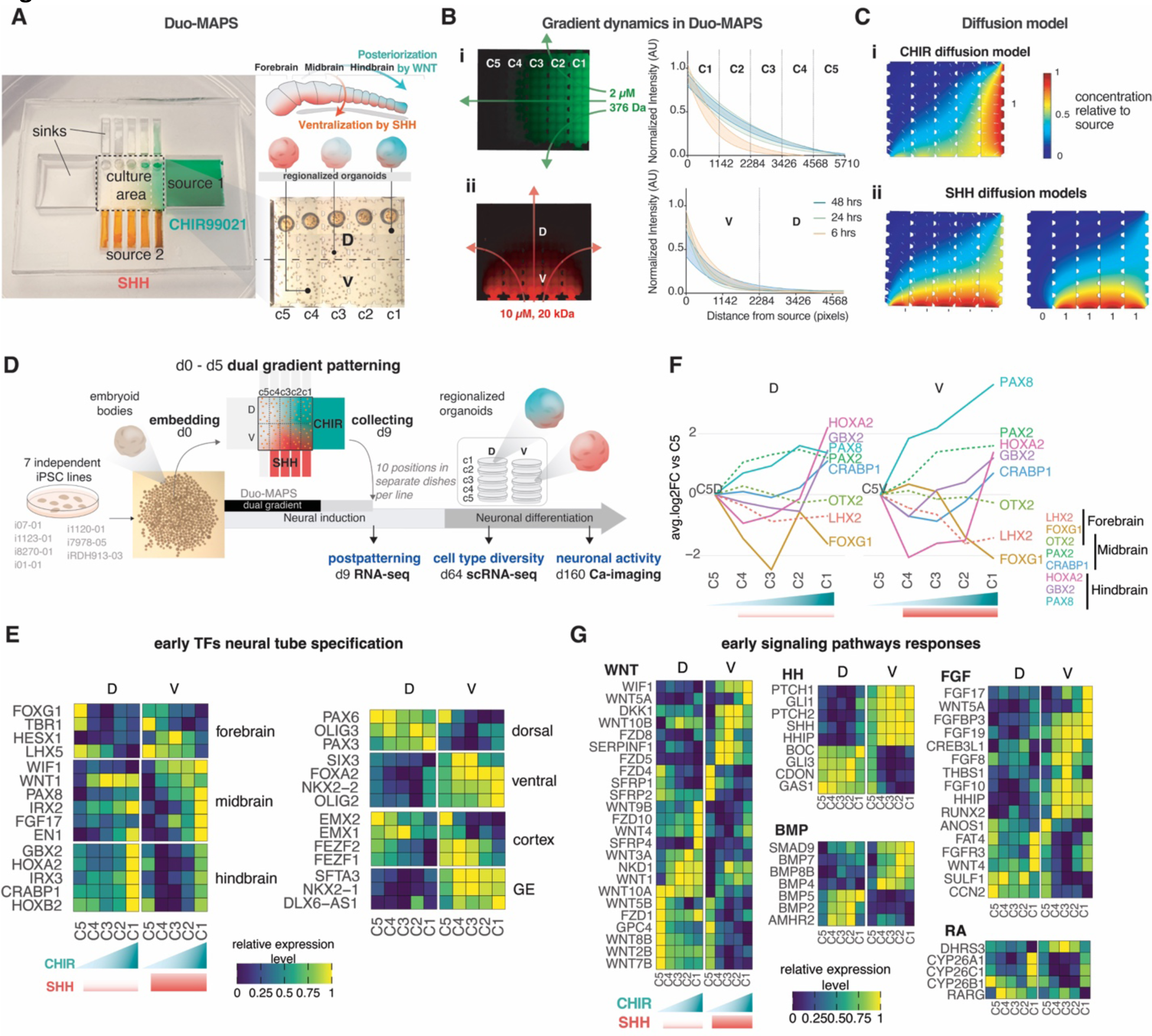
Regional specification of brain organoids by WNT- and SHH-activating gradients in Duo-MAPS. (**A**) Model of the neural tube and schematic picture of the Dual Orthogonal-Morphogens Assisted Patterning System, a mesofluidic device designed to expose organoids to orthogonal morphogen gradients to mimic brain regional specification. EBs are injected in the culture area (bottom right) through pinholes (circles) embedded in hydrogel that once polymerized set them in positions across the culture area. (**B**) Shape, stability, and dynamics of diffusion profiles across the culture area of the device over 48h for fluorescent probes of equivalent molecular weight as CHIR (fluorescein sodium salt, horizontal axis) and SHH (Antonia Red-dextran, vertical axis). (**C**) Simulated diffusion patterns at 24h for 2µM of CHIR along the horizontal axis (top panel), and for SHH along the vertical axis (bottom panels; diffusions with 10 µM of SHH from the bottom (left) and with 100 nM of SHH from the bottom except underneath C5 (right), corresponding to the experimental condition). White arrows represent diffusion vectors. *See also Figure S1*. (**D**) Overview of the experimental design and sample collection to assess the effects of a dual CHIR/SHH gradients. *See also Figure S2*. **E, G)** Heatmap of normalized expression levels across device positions at d9 for genes differentially expressed across positions and known to be involved in brain patterning (**E**) and for genes belonging to the HH, WNT, BMP, FGFR, and RA KEGG pathways (**G**). (GE=ganglionic eminences)(all genes meet ANOVA-FDR < 0.05). **(F)** Line plots showing average log2-fold change (avg.log2FC) relative to the C5 position at d9 along device positions for major forebrain, midbrain and hindbrain markers (significance: continuous lines if ANOVA-FDR < 0.05, dashed lines if FDR > 0.05).

To recreate both antero-posterior and dorso-ventral patterning, we sought to simultaneously generate a gradient of the small molecule CHIR (CHIR99021, a GSK3 inhibitor) along the horizontal axis and a gradient of the protein SHH along the vertical axis, triggering a graded concomitant activation of both the WNT and SHH signaling pathways (**Fig. 1A**). To assess the shape and stability of the diffusion patterns of CHIR and SHH across the culture area, two fluorescent molecules of equivalent molecular weights (374 Da and 20 kDa, respectively) were placed in the source reservoirs and imaged over 48 hours. Analysis of the fluorescence levels suggested that daily replacements of the reservoir’s content ensured stable gradients of CHIR and SHH along orthogonal dimensions (**Fig. 1B**). A simulation of these gradients (**Fig. 1C, Fig. S1, Methods**) revealed a parabolic diffusion pattern for CHIR due to the presence of the five pairs of smaller reservoirs along the horizontal axis, serving as additional sinks for that molecule (**Fig. S1B-D**). The diffusion pattern of SHH along the vertical axis was also parabolic and could be modulated by tuning its concentration in each of its source reservoirs (**Fig.1C**, **Fig. S1Eii**).

We applied this CHIR-SHH dual patterning protocol to 7 human iPSC lines derived from neurotypical male donors (**Table S1**) ^11,21^. EBs were exposed for 5 days (patterning phase, day 0 to day 5) to orthogonal gradients of CHIR (2 µM/mL at the source) and SHH (100 nM/mL at the source) in Duo-MAPS while undergoing an established dual-SMAD inhibition (i.e., BMP and TGFß inhibition) neural induction protocol (**Fig. 1D**, **Fig. S2A-B**, **Methods**). SHH was omitted from the reservoir of the C5 position to model the most anterior region of the neural tube not reached by the notochord *in vivo* ^22^ (**Fig. 1Cii**). At day 9, organoids were harvested from the Duo-MAPS by separating the culture area into a grid of 10 distinct culture positions representing different CHIR/SHH exposure (positions C1 to C5 along the horizontal axis, each separated into D and V portion along the vertical axis, making “C1V” the position exposed to the highest CHIR and SHH concentrations, **Fig. 1A,D**). Patterned organoids were either processed for bulk RNA-seq analysis at d9 or placed in separate dishes to continue the neuronal differentiation protocol and obtain mature organoids from each device position, which were analyzed by single-cell RNA-seq 2 months after patterning (d64 scRNA-seq) and neuronal activity by calcium imaging 3 months after (d170, 5-month-old organoids).

Following patterning at d9, many TFs known to be expressed in specific regions of the neural tube exhibited a graded expression across the culture area’s positions, suggesting that EBs were specified into different brain regions along the gradients (**Fig. 1E-F, Table S2**). Anterior and forebrain markers such as *FOXG1, EMX2*, *LHX5* or *TBR1* showed higher expression in samples from C5-C3; midbrain markers such as *EN1, WNT1* and *PAX2/8* in C3-C1; and hindbrain markers such as *HOXA2/B2* and *GBX2* in C1-C2 (**Fig. 1E-F)**. This antero-posterior specification along the C5 to C1 horizontal axis was achieved both in the D and V portion of the culture area (**Fig. 1F**). Simultaneously, dorso-ventral specification along the vertical axis of the device was demonstrated by higher expression of ventral determinants such as *FOXA2*, *SIX3,* and *NKX2-1/2* in positions closer to the SHH source (C4V to C1V). This included determinants of subcortical structures like the ganglionic eminences, such as SFTA3, LHX6, and DLX6-AS1 (**Fig.1E, F; Fig. S2D**). This ventralization was less pronounced in position C5V where SHH was not added. In contrast, samples located farther away from the SHH source (i.e., C1D-C5D samples) overexpressed TFs such as *PAX3/6, OLIG3,* and *GLI3*, which are associated with dorsal fates and known to be repressed by SHH ^23,24^ (**Fig.1E, F**). To verify that the graded antero-posterior positional identity acquired by EBs was a direct response to differential levels of WNT pathway activation, we compared two unilateral CHIR gradients (2 µM or 4 µM at the source, without any SHH) and showed by RT-qPCR of *GBX2*, *OTX2* and *FOXG1* that a steeper CHIR gradient shifted the specification of EBs posteriorly across the culture area (**Fig. S2F**). We also observed that while CHIR alone could influence genes associated with dorso-ventral patterning, the dual action of CHIR and SHH created a wider range and more topologically accurate expression of those genes in the culture area (**Fig. S2G,H**). Together, these results demonstrated that the dual WNT/SHH gradient was sufficient to trigger in EBs the transcriptional signatures of antero-posterior and dorsal-ventral patterning of the embryonic neuroepithelium.

To understand how this specification occurred, we investigated the expression patterns of genes related to major signal transduction pathways, including the hedgehog (HH), WNT, BMP, FGF receptor (FGFR), and retinoic acid (RA) pathways (**Fig. 1G**). EBs closer to the SHH source (SHH^high^, i.e., C1V-C4V positions) responded to the gradients by producing the *SHH* transcript itself, along with its receptors *PTCH1/2,* the secreted inhibitor *HHIP*^25^ and the SHH direct target *GLI1*. In contrast, EBs exposed to lower SHH concentrations (SHH^low^, i.e., C1D-C5D and C5V positions) showed higher expression of *GLI3*, an inhibitor of the pathway expressed *in vivo* in the dorsal ventricular zone^23^, as well as other SHH co-receptors, like *GAS1*, *CDO*, *BOC* ^26^. The *GLI1/GLI3* inverted pattern confirmed that EBs were able to respond to SHH ^27^. WNT ligands, receptors, and modulators showed more complex expression patterns across the culture area, with peak expression levels either closest to the CHIR source, closest to the SHH source, or furthest to the CHIR source (including the WNT antagonists *SFRP1/2*). These patterns suggested both positive and negative regulations by both CHIR and SHH on different transcripts of the WNT pathway and mimicked in several cases those found *in vivo* ^28-30^. BMP signaling transcripts showed overall lower levels in anteriorized positions where FOXG1 expression was the highest, in accordance with their antagonistic reciprocal regulation during neural tube development^31^. Interestingly, SHH seemed to repress *BMP5/2* and trigger *BMP4/7/8B* expressions ^32^. FGF signaling transcripts were activated mostly in SHH^high^ positions, creating a coordinated expression of SHH and FGF8 known to specify ventral midbrain and hindbrain ^33^. FGF17 and FGF19, which marks the midbrain-hindbrain junction *in* vivo and have been shown to induce cerebellar tissue ^13^, were strongly co-expressed in C1V. The expression of FGF8 in C5D mimicked the presence of the anterior commissural plate ^34^, while FGFR3 was more expressed closest to the CHIR source. Finally, the strongly posteriorizing RA signaling pathway ^5^ was activated under high WNT and SHH conditions (C1 position) as shown by higher levels of the negative regulators CYP26A1/C1/B1.

To rule out any interference of the embedding matrix or other Duo-MAPS components in the response to morphogens, we compared gene expression obtained in EBs exposed to different homogeneous concentrations of CHIR and/or SHH added to the culture medium, including a condition where the WNT pathway was inhibited using the small molecule XAV commonly used to drive organoids towards forebrain fates ^35^ (**Fig. S2D, Methods)**. Organoids patterned in extreme positions of the device revealed similar transcriptomic changes to those observed when morphogens were added to the medium at equivalent concentrations, confirming that the Duo-MAPS approach did not alter the morphogens’ effects (**Fig. S2D-E**). While some genes showed intermediate levels of expression across positions in the device when compared to the extreme conditions tested, many genes (e.g*., NEUROG1, OLIG3, ZIC1/4,*

*LHX9, FGF17*) showed a higher response to intermediate than to extreme doses of morphogen, validating the relevance of our approach to expose EBs to a continuous concentration gradient. Of note, the highest level of *FOXG1* expression was obtained in the presence of XAV, consistent with data suggesting that WNT inhibitors secreted from the anterior mesoderm are involved in forebrain specification during gastrulation ^36,37^. However, this was not the case for *FEZF2*, *EMX2*, *PAX6* or *TBR1*, which had similar expression in XAV, basal (dual-SMAD only), and C5D. Finally, we used this dataset to identify the sources of variation driving gene expression across all samples at d9 ^38^, delineating the contribution of both morphogens and line-to-line variations, which suggested that while the morphogens triggered strong changes in gene expression, some genes were also susceptible to show differences across iPSC lines (**Fig. S2I, Table S2**).

Overall, results showed Duo-MAPS’ ability to expose EBs to orthogonal gradients of CHIR and SHH created a cascade of intracellular signaling and transcriptomic changes along the culture area, efficiently recreating the antero-posterior and dorso-ventral axes associated with early neural tube patterning. Difference of results for different graded SHH and CHIR concentrations underscores the importance of studying 2D rather 1D gradients of simultaneously imposed morphogens to model *in vivo* brain development.

### 2. Patterned organoids generated the major neuronal lineages of the human fetal brain

We characterized the cell types generated by organoids derived from 6 iPSC lines by scRNA-seq 2 months after Duo-MAPS patterning (d64, multiplexed data from 2 experimental batches, **Methods**). Cells from all samples were integrated into a common dataset (198,405 cells in total) and clustered by transcriptome similarity ^39^. The obtained clusters were annotated based on specific gene expression (**Table S3, S4, S5**, **Methods**) and grouped in 16 major cell types or lineages, associating them - when possible - with corresponding putative brain regions based upon known gene markers (**Fig 2A-C, Fig. S3A**). In UMAP projection, cells could be broadly separated into different progenitor cells, including radial glial cells (RG) and proliferating progenitors, intermediate progenitor cells (IPC), and immature and mature neuronal cells (**Fig 2A, C**). Neuronal cells branched into multiple subtypes, including *GAD1/GAD2/SLC32A1(vGAT)+* GABAergic neurons and *SLC17A6*(*vGluT2*)+ or *SLC17A7*(*vGluT1*)+ glutamatergic neurons (**Fig. 2B-C, Fig.S3B**). RNA velocity analysis followed by latent time inference ^40,41^ confirmed the transcriptomic progression of differentiation in captured cells along each of those lineages, from progenitors to mature neurons (**Fig.S3C**).

**Figure 2:**
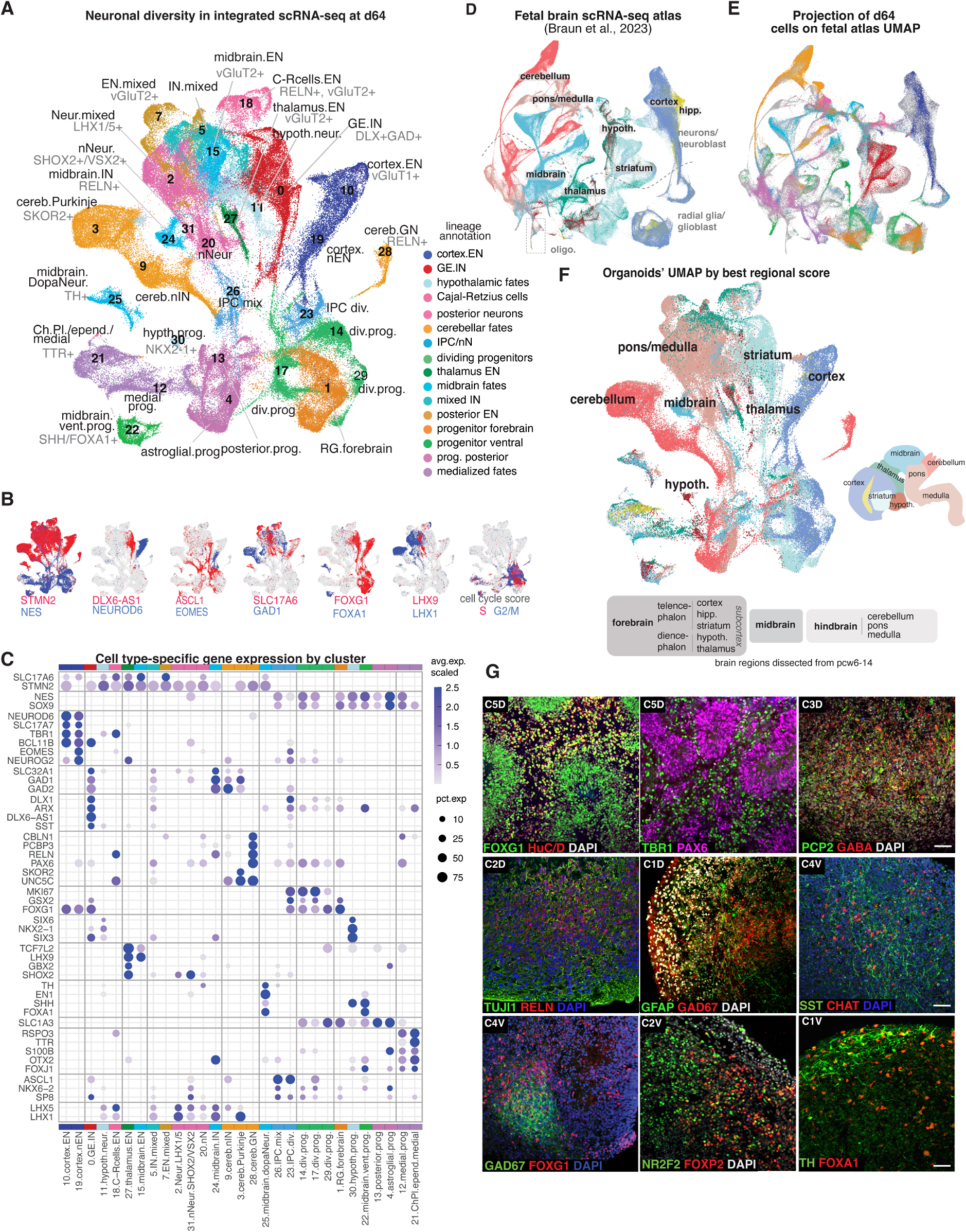
Diversity of cell type generated in d64 organoid reproduce multiple fetal brain lineages. **A)** UMAP plot of integrated scRNA-seq d64 organoids with cells colored by major lineage/cell types with each cluster’s number, annotation, and defining genes. **B)** UMAP plots showing expression levels for pairs of contrasting genes (blue vs red, with limited co-expression in purple). Rightmost plot: cell cycle score for S and G2/M phases (**Methods**). **C)** Dot plot of average expression values per cluster (avg.exp.scaled) for known marker genes defining cell types and/or regional identities. Dot size shows percent of cells expressing the gene (pct.exp > 5%). **D)** UMAP plots of scRNA-seq integrated data from a human fetal brain atlas (reprocessed from Braun et al, 2023) ^42^ colored by dissected region of origin, as indicated by schematics (hypoth.=hypothalamus, hipp.=hippocampus, oligo.=oligodendrocyte precursors). **E)** Projection of d64 organoid cells (colored as in A) onto the fetal brain UMAP based on transcriptomic similarities (see **Methods**) confirming the correspondence between organoids and brain regional lineages. **F)** Organoid UMAP colored by highest predicted region of origin following projection on the fetal brain reference (predicted score > 0.4). **G)** Representative immunofluorescence images for protein markers of some of the regions and cell types present in d64 organoids, including forebrain (FOXG1), cerebellar Purkinje cells (PCP2); Cajal-Retzius cells (RELN), GABAergic neurons (GAD1), astrocytes (GFAP); cholinergic neurons (CHAT), MGE neurons (SST), CGE neurons (NR2F2), LGE neurons (FOXP2), and dopaminergic neurons (TH/FOXA1). Organoids’ positions of origin in the device indicated (e.g. “C5D”). scale bar = 50 µM.

To inform and validate our cluster annotation of cell type and brain regional identity, we integrated our dataset with a recently published scRNA-seq atlas of the fetal human brain ^42^, which collected a broad range of regions during the first trimester of gestation (i.e., postconceptional week 6-14) allowing us to compare organoid cells to *in vivo* cell types (**Fig. 2D, Fig. S4**). We first projected organoid cells on this reference atlas using UMAP dimensionality reduction-based projection and label transfer methods from *Seurat* to label organoid cells with the closest corresponding fetal brain (**Methods**). Cells from patterned organoids matched the transcriptional signatures of progenitors and neurons from the fetal cortex, striatum, thalamus, and hypothalamus as well as from the midbrain, cerebellum, and pons/hindbrain, with more limited match for hippocampus and medulla (**Fig. 2D-F**). In many cases, organoids reproduced the distinctive gene expression patterns exhibited by lineages and regions of the brain (**Fig.S3D-E, Fig. S4**). This regionalization was also confirmed using the Voxhunt tool ^43^ by projecting organoid clusters on the different regions of the E15 mouse fetal brain (**Fig. S3F**).

Forebrain lineages marked by FOXG1 included a cortical-specific excitatory lineage (19.cortex.nEN, 10.cortex.EN *BCL11B/NEUROD6/SLC17A7*+) and a subcortical-specific inhibitory lineage related to fates of the ganglionic eminences (0.GE.IN, *DLX1/2/5/6+*). In the latter, we could identify cells related to the medial (*SST*/*NKX2-1/LHX6*+), caudal (*PAX6/CALB2/SCGN*+), and lateral (*SIX3*/*ZNF503/EBF1/MEIS2+)* ganglionic eminences fates, including cells expressing markers of medium spiny neurons’ precursors ^44^ (**Fig. S4C, Fig. S5A-C**). Organoids also contained Cajal-Retzius cell-like excitatory neurons (18.C-Rcells.EN, *RELN/SCL17A6/TP73/TBR1+*), an early neuron of the cortical preplate^35^. Diencephalon fates included two clusters corresponding to ventral progenitors and neurons of the hypothalamus (30.hypth.prog., *SIX6*/*DLK1/RAX/NKX2-1/SHH+,* and 11.hypoth.neur., **Fig. S3A,E**), and a cluster of excitatory neurons characteristic of the thalamus (27.thalamus.EN, *GBX2/TCF7L2/LHX9/SHOX2/SLC17A6+*) (**Fig. 2A,C, Fig. S4F**). In addition to anterior fates, a progenitor branch presented a signature of medialized fates corresponding to the choroid plexus, cortical hem, hippocampus, and/or rhombic lip in the posterior brain (clusters 12 and 21, *TTR/RSPO3+*) (**Fig. 2C**). As for the choroid plexus *in vivo*, 50% of the cells in cluster 21 expressed the serotonin receptor *HTR2C* which is involved in the formation of the cerebrospinal fluid ^45^ (**Fig. S3A**).

Four distinctive clusters showed midbrain-like signature, including excitatory neurons (15.midbrain.EN, *TCF7L2/LHX9/SCL17A6+,* and *GBX2*-negative), inhibitory neurons (24.midbrain.IN, *RELN/OTX2/TAL1/TAL2/GATA2/GATA3+*), a ventral progenitor cluster (22.midbrain.vent.prog., *FOXA1/LMX1A/SHH/ARX+*), and a related dopaminergic neuron cluster (25.midbrain.dopa.neur.) (**Fig. S4D-G, Fig. S3E**)^46^. The dopaminergic neuron cluster expressed canonical dopaminergic markers (*TH*/*NR4A2*(*Nurr1*)/*DDC/PITX3*/*EN1/EN2+)* but no noradrenergic or serotoninergic markers ^47^. Fates from the cerebellum included a Purkinje cell cluster (3.cereb.Purkinje, *PCP4/SKOR2/GAD1/CA8/UNC5C+*), a GABAergic immature interneuron cluster (9.cereb.nIN, *PAX2+*), and a smaller cluster of granule neuron (28.cereb.GN, *PAX6/RELN/PCBP3/CBLN1+*), in line with the later specification of granule neurons *in vivo*^48^. All three clusters had strong transcriptomic correspondence to the cell types described in a human fetal cerebellum scRNA-seq study ^49^ (**Fig. S5D-G**). Consistent with their distinct origins in the cerebellar primordium, immature interneurons expressed the ventricular zone marker *PTF1A* and granule neurons expressed the rhombic lip marker *ATOH1* (**Fig. S5H, Fig. 2A,C**).

Progenitor cells with mixed posterior signature between hindbrain and midbrain lineages (13.posterior.progenitors, 4.astroglial.progenitors, 26.IPC.mix, 20.nN) generated posterior neuronal lineages with less defined identities, including inhibitory neurons (5.IN.mixed) and excitatory neurons (7.EN.mixed cells) matching cells from multiple regions (i.e., midbrain, pons, thalamus), as well as posterior clusters (2 and 31) expressing *LHX1*, *LHX5* or *SHOX2,* TFs expressed *in vivo* from the diencephalon to the pons ^50^ (**Fig. S3A,E**, **Fig. S4D**). Mixed regional signatures suggest that cells were too immature to be confidently identified -organoids generating less distinguishable cell types than *in vivo* ^51^ - or represented lineages shared or migrating between brain regions. At the used morphogen concentrations, only limited expression for posterior hindbrain/spinal cord or neural crest fates was evident at d64.

Immunofluorescence colocalization analysis confirmed the presence of FOXG1+ telencephalic lineages, TBR1/PAX6+ cortical neurons, SST+ medial ganglionic eminence interneurons, NR2F2/FOXP2+ caudal ganglionic eminences neurons, PCP2(Purkinje cell protein-2)/GABA+ Purkinje cells, RELN+ C-R cells and TH/FOXA1+ dopaminergic lineages (**Fig. 2G**). While cholinergic markers (*CHAT*, *SLC18A3*/*SLC5A7*) were sparsely present in scRNA-seq data, the presence CHAT+ cells suggested the production of cholinergic neurons (**Fig. 2G**).

In conclusion, patterned organoids generated the major transcriptomic lineages observed across many regions of the early fetal brain.

### 3. Cell lineages generated in organoids were influenced by early morphogens exposure and by iPSC line

To understand how exposure to the WNT agonist and SHH gradients influenced the diversity of cell fates observed in patterned organoids across iPSC lines, we performed a detailed analysis of cell composition across the 60 scRNA-seq samples collected at d64 (10 positions x 6 genotypes, **Fig. 3**, **Fig. S6**). Most cell types were produced by organoids from multiple device positions with stronger differences in cell types between D and V portions of the device than across the C1 to C5 positions (**Fig. 3A**,**B****, Fig. S6A,B**). In addition, while all cell types were observed in organoids from all 6 iPSC lines tested, the relative proportions of cell types present at each device position varied across individuals (**Fig. 3A,F,G, Fig. S6C-E**). Variance analysis confirmed that the influence on cell proportions overall was driven by differences between D/V positions, difference between genotypes, and difference between C1 to C5 positions, in that order (permutational MANOVA R2 = 0.244, 0.190, and 0.154, respectively, all with p-values < 0.001). Principal component analysis showed that this variation was driven by the differential coproduction of certain cell type groups between organoid samples (**Fig.S6F**).

**Figure 3.**
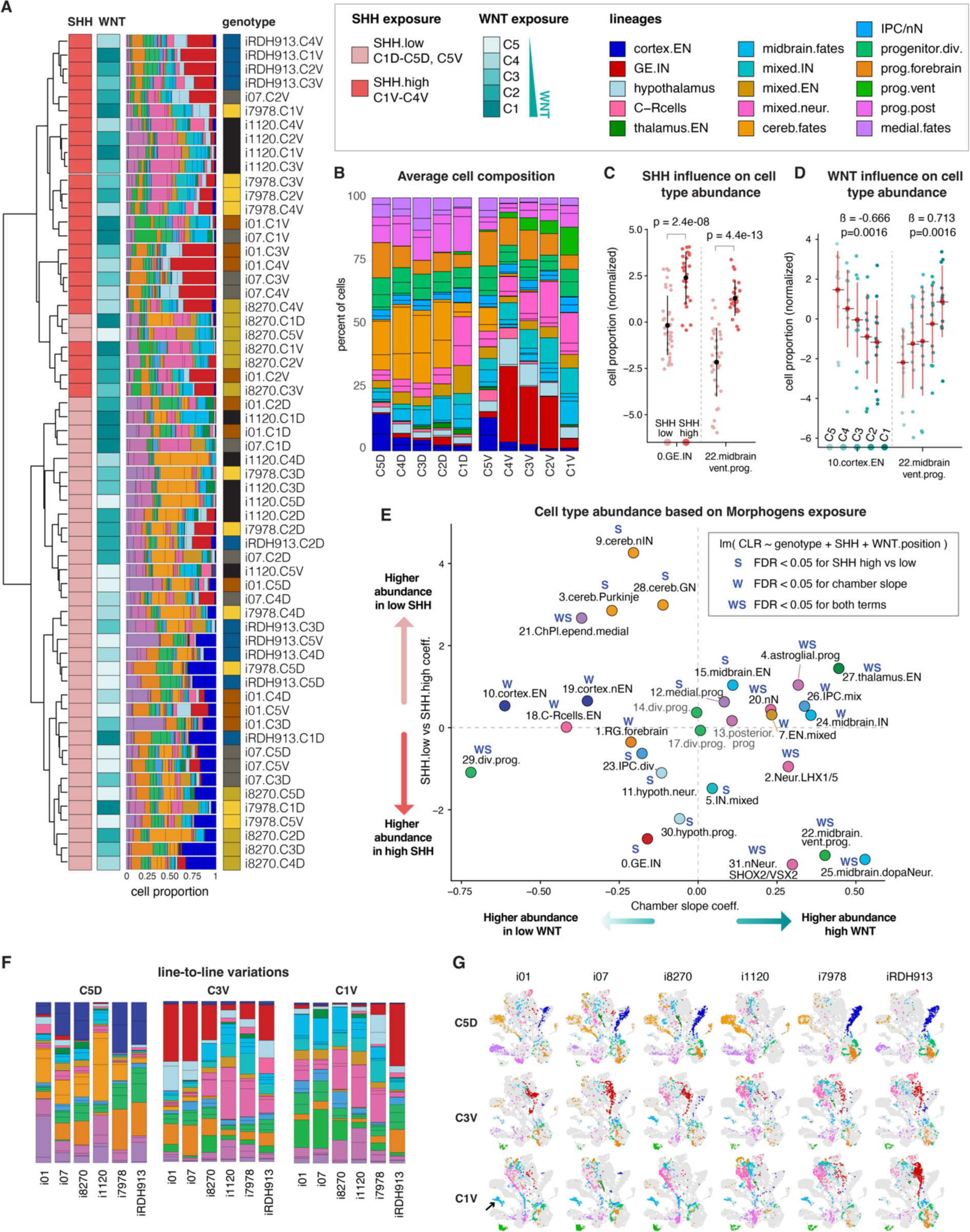
Cell lineage production in organoids was influenced by early morphogen exposure. **A.** Hierarchical clustering of organoid samples’ cell composition with corresponding cell composition bar plot colored by main identified lineages. Samples annotated based on genotype, WNT agonist and SHH exposures (see **Table S4**) **B.** Bar plot of averaged cell type composition across Duo-MAPS culture positions (n=6 lines per position). Color code as in **A**. **C.** Dot plots for two clusters showing significant differences in cell proportions between samples from SHH^low^ and SHH^high^ (p-value: two-sided t-test on normalized proportions, mean and standard deviation shown in black). See also **Fig. S6F**. **D.** Dot plots showing significant linear trend of cell proportion changes across positions along the WNT gradient (from high WNT =C1 to low WNT=C5, shades of blue). Slope and p-value by linear model shown above, mean and standard deviation shown in red. **E.** Dot plot showing the inferred influence of SHH (SHH^high^ vs SHH^low^ coefficient (coeff.), y-axis) and WNT exposure (C1 to C5 slope, x-axis) on the cell proportion of each cluster. Linear model-based significant cases for either or both coefficients indicated by “W”, “S”, “WS” (see box). **F.** Bar plot of cell composition showing example of variation in cell proportions across lines at 3 specific positions. **G.** Illustrative UMAP plots by lines and positions as **F** (samples from each line subsampled to same number of cells).

We then used linear models to evaluate the overall influence of SHH (SHH^high^ vs SHH^low^) and WNT gradients (C1 to C5 slope) on cell type production (**Fig. 3C-E Methods**). Organoids exposed to high SHH conditions generated significantly more ganglionic eminence and hypothalamic progenitors/neurons (clusters 0, 11, 30), while organoids exposed to low SHH conditions generated significantly more cerebellar fates (clusters 3, 9, 28). Organoids exposed to lower levels of WNT generated higher fractions of forebrain radial glia, cortical excitatory neurons, and Cajal-Retzius (C-R) cells (clusters 10, 19, 18, 1). Organoids exposed to both high SHH and high WNT generated higher fraction of midbrain dopaminergic neurons and ventral progenitors (clusters 25, 22), as well as pons and hindbrain lineages (clusters 31, 2). Finally, clusters generated under high WNT but low SHH conditions included thalamic excitatory neurons (cluster 27), suggesting that their production was repressed by SHH exposure. Overall, regional fates required different ranges of CHIR/SHH concentrations to be generated efficiently.

These changes in cell proportion marking SHH-driven ventralization and CHIR-driven posteriorization were also supported by differential gene expression analyses between device positions in progenitor cells at d64 (D vs V, and C1 vs C4/C5 DEGs in cluster 1, 4 and 13, **Fig. S7A-B, Table S5**). Progenitors expressed higher levels of dorsal-associated TFs in samples from low SHH conditions and higher level of ventral-associated TFs in high SHH conditions across most iPSC lines (e.g., *LHX2, PAX6, EMX2, PAX3, GLI3 vs NKX6-2, DLX2, GSX2, SIX3, FOXP2*). Similarly, comparing position C1 vs. C4/5, progenitors upregulated posterior genes such as *IRX3/5, NKX6-2 and FOXB1,* and downregulated anterior genes such as *FOXG1, SOX6,* and *LHX2*, although this was less robust across lines.

The different distributions of cell types across positions were also validated by differential protein distribution. Immunostaining confirmed the predominance of PAX6-positive cells in organoids patterned with low SHH (D positions), while NKX2-1-positive cells were predominant in high SHH (C4V to C2V positions, **Fig. S7C-F**). Similarly, decrease in the percentage of TBR1/PAX6+ cells between C5D and C1D positions, with some line-to-line variation, validated the WNT-induced posteriorization quantified by scRNA-seq (n=4 individuals**, Fig. S7G-I**). High performance liquid chromatographic-coulometric (HPLC-EC) analysis at day 78 revealed the presence of dopamine (DA) detected in a range of 23 to 820 pg per single organoid tested depending on organoid position in the device, with higher dopamine levels in organoids from C1V and progressively decreasing levels in samples C3V to reach the lowest levels in C5D (**Fig. S6G**). The presence of serotonin (5-hydroxytryptamine, 5-HT), produced by neurons in the ventral region of the hindbrain, was detected in the two lines in C1V, confirming the generation of hindbrain-like cells in this position of the device, while minimal 5-HT levels were detected in C5D-derived organoids (<10 pg/organoid).

All analyses of cell composition, differential expression and protein distribution pointed to a consequent cell line-specific effect on the results, confounding in parts the overall effects of SHH and CHIR exposure (**Fig.3A,F-G Fig.S6C-G**). For instance, there were noticeable differences in the number of dopaminergic neurons produced in C1V across individuals (TH+cells, 25.dopa neurons, **Fig.S6C-E**). Similarly, a notably high level of serotonin was identified in C3V-derived samples from the i8270 line (327 pg/organoid), while C3V-derived organoids from the i01-01 line displayed a considerably lower 5-HT level (23 pg/organoid) (**Fig. S6G**), supporting the increased proportion of hindbrain cells observed in i8270 C3V-derived samples (clusters 2.Neur.LHX1/5, c31.nNeurSHOX2/VSX2 in **Fig. S6C,E**). Even lineage that were strongly influenced by morphogens, such as GE IN or cortex EN, show certain degrees of variation between lines (**Fig.S6E, S7**). These observations suggested that the different lines exhibited different responses to morphogens and/or different propensity to generate identical lineages under the same morphogenic conditions.

Altogether, these analyses revealed the complex influence of early exposure levels of WNT and SHH on the production and specification of neuronal lineages. They also highlight that iPSC lines tended to respond differently to the same morphogenic conditions, warranting a deeper analysis on how transcriptomic profiles established at d9 controlled neuronal specification across individual genotypes.

### 4. Neuronal lineages arose from early programs, generated by crosstalk between dual morphogen signaling in precursor cells

The Duo-MAPS offers a unique perspective to identify the transcriptional programs triggered in response to the combined effects of SHH and CHIR concentration gradients in different iPSC lines. To identify their overall effects and extend on our initial characterization (**Fig. 1E-F**), we performed unbiased clustering of all genes differentially expressed at d9 across the culture area to identify 20 main *morphogen-response spatial modules (*p1 to p20, k-mean clustering on normalized average expression, **Fig. 4A, Table S2**). Those spatial modules decomposed the gene expression programs triggered under specific levels of SHH and WNT pathways activations as shown by their complex spatial expression trends across the culture area (**Fig. 4A**). Gene expression triggered at the lowest levels of a morphogen could indicate an overall repressive effect of the morphogen on the genes belonging to the modules (e.g., p1-3 indicated repression by WNT and SHH). Similarly, genes showing linear trends across either the long or short axis could be interpreted as solely dependent on WNT or SHH level, respectively (e.g., p10 or p19), whereas genes showing bell-shaped pattern along the horizontal axis suggested a combination of SHH activation and WNT repression (for instance p17 or p13). Due to our sampling approach of the culture area (i.e., 5x2 positions) WNT-response signal was better decomposed than the binary SHH-response (activated/repressed). The activation of those morphogen-response patterns early on was often correlated with the production of neuronal lineages in organoids later (**Fig. 4B, Methods**). Finally, the authenticity of those WNT/SHH-response patterns established using our approach was assessed *in vivo* by evaluating their expression in progenitor cells from the 1^st^ trimester brain scRNA-seq atlas (**Fig. 4C,D, S8A-B, Method**).

**Figure 4:**
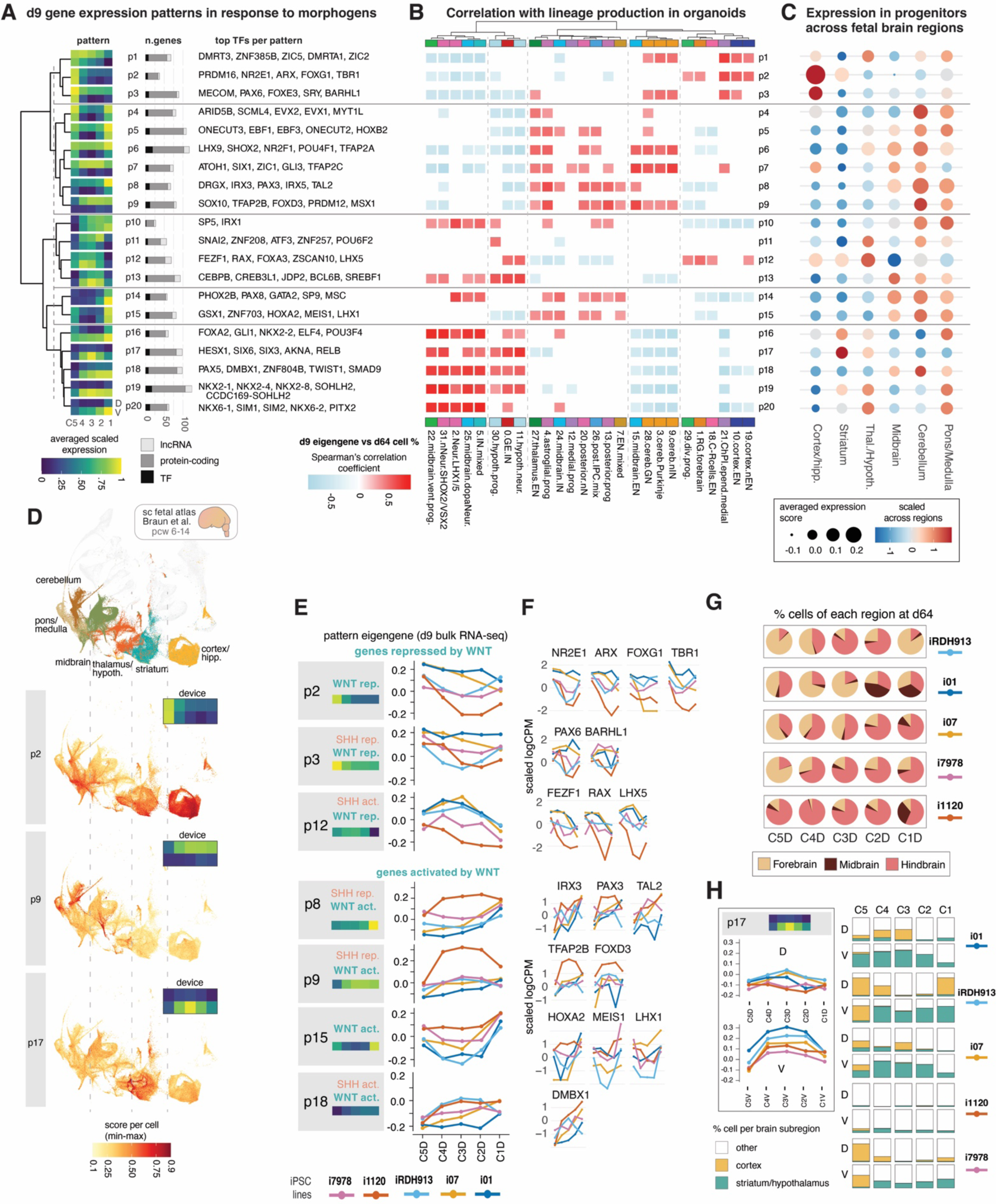
Early SHH and WNT gradient exposure-response association with late neuronal cells generation and variation between iPSC lines. **A)** Clustering of morphogen-response expression pattern at d9 (k-mean on min-max normalized expression, named “p1” to “p20”). Only genes with significant differential expression across Duo-MAPS positions were clustered (ANOVA-FDR < 0.05). Corresponding averaged expression trend across the culture area for each pattern plotted as separate heatmap. Patterns were organized by hierarchical clustering (Ward’s method). Corresponding bar plots show number and type of transcripts per pattern and the Top TFs ranked by FDR in each pattern are listed (full gene list in **Table S3**). **B)** Heatmap of correlation between the eigengene of d9 expression patterns defined in A and cell type proportions at d64 (Spearman’s correlation coefficient on n=51 samples with matched d9 bulk RNA-seq log2CPM and d64 scRNA-seq cell proportion data, **Table S1,** two-sided FDR-corrected p-value < 0.05). **C)** Dot plot showing average expression score of each pattern in progenitors from each major region of the brain atlas ^42^ (dot size). Dot colors shows scaled values of expression scores normalized across region (expression score was computed using Seurat *ModuleScore* function using all genes included in each pattern, **Methods**). **D)** UMAP plots showing the corresponding expression score of 3 morphogen-response patterns from **A** in progenitor cells from the fetal brain atlas, (region of origin shown by color on the top UMAP). Score was min-max normalized across all cells. The corresponding expression pattern in the device given as in **A** for comparison. **E)** Left panel, average expression trend across the culture area for pattern of genes in D position (C1D-C5D) responding to WNT/SHH (act.=activated, rep.=repressed). Right panel, line plots of eigengenes levels at day 9 for iPSC lines from 5 individuals (colors) for the spatial modules shown on the left(see also **Fig. S8C**). **F)** Line plots showing normalized expression values across lines at d9 of genes selected from each pattern in panel **E** (scaled log2CPM). Note the outlier response displayed by line i1120. **G)** Pie chart representing the fraction of cells falling into the 3 major antero-posterior brain subdivisions at d64 (from fetal brain atlas projection) separated by lines and device positions. Note the more pronounced posterization of line i1120. **H)** Spatial module p17 eigengene at d9 in V positions shows enhanced response to SHH of i01 and RDH913 iPSC lines as compared to other lines. Bar plots showing the percentage of cells associated to the fetal brain striatum and/or hypothalamus compared to cortex and other fates across lines and device positions.

To illustrate, the gene programs repressed by both SHH and WNT at d9 (i.e., p1, p2, p3, **Fig. 4A**) and associated with cortical EN production in organoids (**Fig. 4B**) involved *FOXG1*, *PAX6, DMRTA1*, *DMRT3*, and *ARX*, known determinants of the dorsalization of the telencephalon. ^52,53^ p2 and p3 were correspondingly highly expressed in progenitors from the fetal cortex (**Fig. 4C**). Gene programs triggered in response to concomitant high SHH and high WNT (i.e., p14, p16, p18, p20) are associated with the production of several midbrain and hindbrain lineages in organoids including known determinants of ventroposterior fates ^54-56^, such as *PHOX2B*, *PAX8*/*5*, *DMBX1*, *NKX6-1/6-2, SIM1/2* or *FOXA2*, with p14, p18 and p20 showing higher expression in posterior progenitors of the fetal brain. Among patterns activated by SHH (p10 to p20), the patterns showing decreased expression in the presence of WNT (i.e., p12, p13, p17) involved TFs such as *RAX*, *SIX3*, and *SIX6*, and were strongly associated with fates of the GE/striatum and hypothalamus in organoids and in the brain for p12 and p17, suggesting that specification of those regions involved sets of genes repressed by WNT exposure *in vivo*^57,58^. Gene patterns repressed by SHH but showing different trends of WNT activity were associated with the production of neurons of the cerebellum (p6,p7,p9). Finally, projecting morphogen-response expression patterns onto progenitor cells from the fetal brain atlas revealed both widespread and restricted expression to specific group of cells, indicating that those cells were generated in vivo under specific dosages of SHH/WNT exposure (e.g, compare p17 with p2 in **Fig. 4D**). This analysis offers a detailed view on how the combination of two morphogenetic signals activating specific transcriptional profiles regulated the regional location of human cells across the brain.

The morphogen-response patterns delineated expression trends averaged across genotypes, identifying gene groups with overall linear increase, linear decrease, or bell-shaped responses to each morphogen. To better understand how each individual line responded to the same gradient, we used the eigengene of morphogen-response modules at day 9 as a measure of the “responsiveness” of individual lines to SHH and WNT (**Fig. 4E**). A different propensity for activating these patterns was observed in each individual iPSC line, which then drove a different cellular composition at d64. Confirming that many genes influenced by WNT exposure drove high interindividual variations (**Fig. S2I, Table S3**), there was an overall higher interindividual variability in the expression of morphogen-responsive patterns along the WNT gradient under low SHH exposure (i.e. C5D to C1D), reflecting a stronger variability in WNT response than in SHH response between individuals (**Fig. 4E**, **Fig. S8C-D**). This differential WNT response resulted in a bias in the specification of cells along the brain antero-posterior axis for the different iPSC lines (**Fig. 4E, F**). Notably, line i1120 showed hyper-responsiveness to the WNT gradient as shown by the increased level of patterns overall activated by WNT (p8, p9 and p15, including *PAX3, FOXD3, TFAP2B and HOXA2*) and lower level of pattern repressed by WNT (p2, p3 or p12) compared to other lines in the D positions of the device (**Fig. 4E,F, Fig. S8C**). This stronger early posteriorization explained the higher proportion of cerebellar and hindbrain-related cells observed in i1120 organoids across all device positions compared to other lines (**Fig. 4G, 3F-G, Fig. S6C,E**). Comparatively, lines i01 and iRDH913 were relatively hypo-responsive to WNT (i.e., lower p8, p9, p15 levels, higher FOXG1 **Fig.4E-F**) and seemed in turn more responsive to SHH as shown by higher levels of p17 (**Fig. 4E,H**). This explained the higher proportion of antero-ventral fates (basal ganglia and hypothalamus) in organoids derived from those lines (**Fig. 4H, Fig. S6C**). The data suggested that different levels of sensitivity to morphogens between individual iPSC lines drove the differences in regional specification and cell composition observed in patterned organoids, attributable to differences in the genetic or epigenetic characteristics of each line, stochastic variation between lines, and/or experimental batches.

### 5. Morphogens activated regulators defining and regulating lineage-specific expression

To better understand how neuronal lineages are specified differently across brain regions, we sought to identify the set of TFs whose expression and regulatory activity defines each cell lineage produced in patterned organoids across the 2 time points.

To measure TF activity in d64 organoids, we generated a gene regulatory network by applying the SCENIC pipeline ^59^ to our scRNA-seq dataset (**Table S3**). SCENIC identified the “regulon” of each TF (i.e. the list of genes putatively activated by the TF) (see **Methods**). The activity of each TF was then inferred from the expression of its regulon across cells to identify the most specific regulators of each cell type (**Fig. S9A-C**). Those cell type-specific regulators included both known and novel combination of TFs (**Fig. S9B-C**). The regulons of identified TFs could be merged to generate regulatory subnetworks driving gene expression specific to each cluster or group of clusters (**Fig. 5A,B, Fig. S9**). For instance, the GE inhibitory neurons subnetwork displayed a positive cross regulatory interaction among the 4 DLX genes and ARX, with secondary interactions with VAX1, FOXG1, MEIS2, and EBF1 (**Fig. 5B, S9A,B**). The subnetwork specific for midbrain ventral progenitors and dopaminergic neurons presented a cross-regulation of FOXA1, FOXA2 and EN1 as a self-reinforcing central triad (**Fig. 5A**).

**Figure 5:**
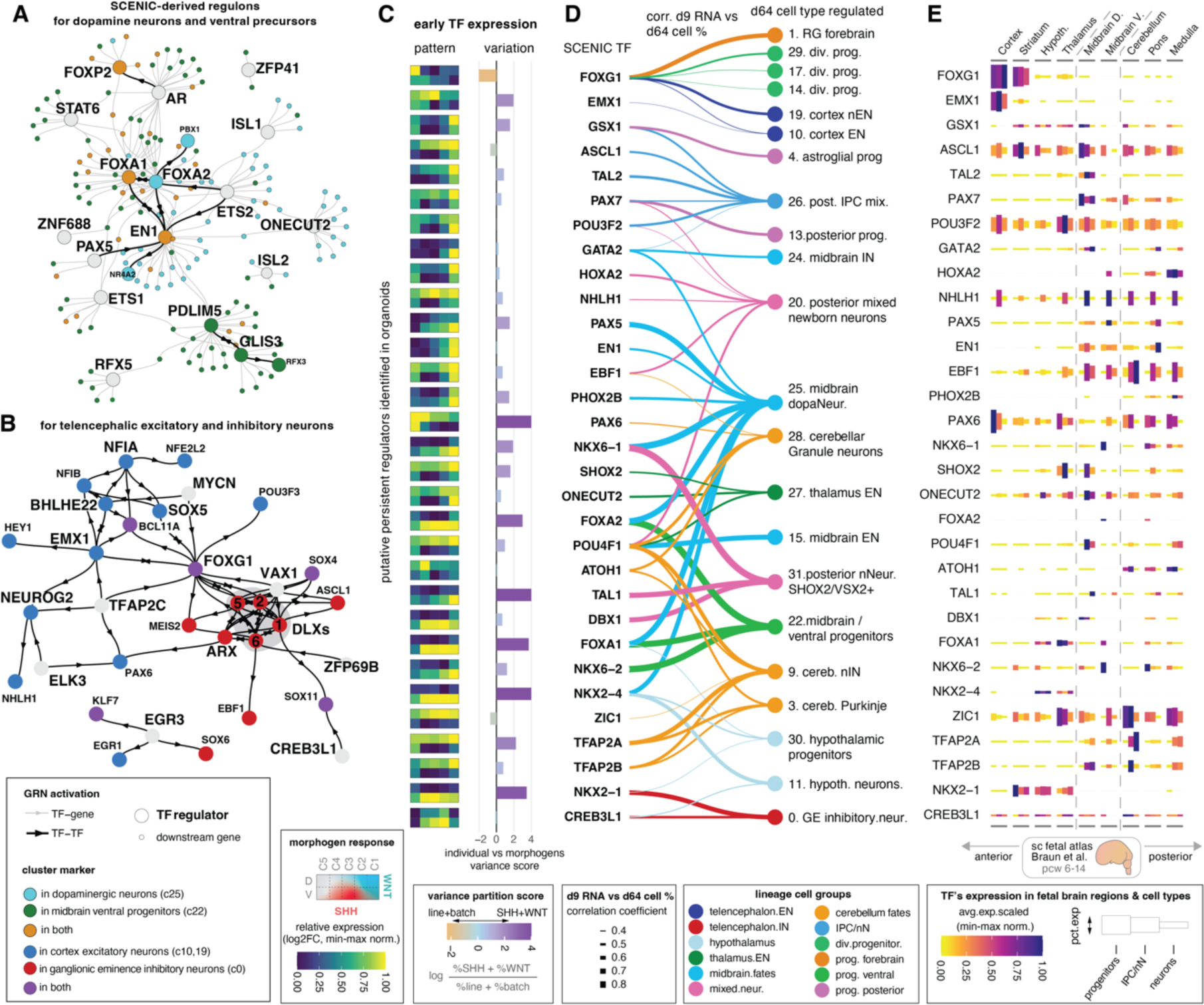
Early regulators with persistent regulatory function during development define brain lineages. **A-B)** SCENIC-derived regulatory subnetworks at d64 focused on TF regulating genes specific to dopaminergic neurons and midbrain ventral progenitors (**A**) or specific to cortical EN and ganglionic eminences IN (**B**). Networks were obtained by merging the regulons of the top 10 specific regulators of selected clusters (ranked by RSS metric, **Table S6**, **Methods**). Dot color indicates genes identified as cluster markers for either or both clusters selected (**Table S5**). Edges are activating and black indicates TF-to-TF regulation (“bowtie” shape indicates bidirectionality). Only TF-TF edges are shown in **B**. *See also **Fig. S9***. **C)**. Expression in the Duo-MAPS at d9 for SCENIC regulators identified in **D)** (bulk RNA-seq, log2FC min-max normalized, ANOVA-FDR < 0.05). Bar plot shows *VariancePartition* results as a log-ratio of the percentage of variation explained by SHH and WNT exposure divided by the percentage of variation explained by line-to-line differences (i.e., interindividual and experimental batch) across all d9 bulk RNA-seq samples collected in this study (**Fig.S2I, Table S2**). **D)** Graph connecting specific regulator TFs identified at d64 by SCENIC with their associated cell type. Association based on 3 criteria: i) TF differentially expressed at d9 (in **C**); ii) TF’s d9 expression significantly correlated with d64 cell percentage across samples (linewidth, FDR < 0.05, **Table S6**); and iii) TF among the top 20 specific regulators for that cell type (by RSS metric in d64 SCENIC regulatory network). **E)** Expression of TF regulators from **D** summarized across the entire fetal brain atlas. Scaled average expression (avg.exp.scaled) computed across major brain region of origin and cell types from the atlas ^42^. Box size proportional to the percent of cells expressing the gene (pct.exp).

We then identified which of cell lineage-specific regulators identified by SCENIC analysis at d64 was also showing an early expression at d9 that correlated with the proportion of that lineage at d64 (correlation coefficient between TF’s d9 RNA level and cell type proportion in d64 across matched samples, **Table S6**). This analysis across time revealed 31 **lineage-defining regulators** with persistent activity across development (**Fig. 5C-D**). For each lineage-defining regulator, we characterized early expression patterns in response to the SHH/WNT dual gradient (**Fig. 5C**), degree of interindividual variations (**Fig. 5C**), strength of the correlation between early expression and cell lineage proportion at d64 (**Fig. 5D**), as well as the TF’s expression level across regions of the human fetal brain atlas (**Fig. 5E**). This multifaceted characterization confirmed that cell lineages were specified early during development, after morphogen exposure, through the activation of canonical TF genes that exert regulatory action on many downstream genes of the lineage. Early expression of those regulators was primarily influenced by SHH-WNT exposure. There were, however, a few notable exceptions, where the genotype had prominent influence on their expression *in vitro* (e.g., FOXG1, ASCL1) (**Fig. 5C**, using variance partition data presented **Table S2**), contributing to the difference we observed between individuals. Many of those cell lineage-defining regulators identified *in vitro* were expressed in the brain region were that cell type is found in the human brain atlas (**Fig. 5E**). For instance, SHOX2 was identified as a master regulator of excitatory neurons of the thalamus and showed enriched expression in progenitors, IPC, and neurons in the thalamus *in vivo*, and to a lesser extent in dorsal midbrain. The regulators of cerebellar interneuron and Purkinje cells TFAP2A and TFAP2B identified in organoids were also expressed in the cerebellum of the human brain and were recently linked to cerebellar interneurons specification in mice ^60^ (**Fig. 5E**, **Fig. S4D**).

### 6. Dorsalized and ventralized organoids displayed distinct spontaneous neuronal activities

To characterize neuronal activity in regionally patterned organoids mimicking the development of distinctive human brain regions we performed functional analysis of sliced organoids from antero-dorsal (i.e., from the C5D position, generating mostly dorsal cortex) and antero-ventral conditions (i.e., from the C4V position, generating mostly basal ganglia and diencephalon) in 3 different lines (**Fig. 6A,B Fig. S10**). We used AAV-mediated expression of the ultrafast protein calcium sensor GCamp8m ^61^ under the *Synapsin* promoter to record calcium concentration changes over time (Ca^++^ transients) as an indicator of neuronal activity. Recordings were performed after 5-months in culture (1-month after slicing), when spontaneous Ca^++^ transients were observable (**Supplemental Methods, Fig. 6C,D, Suppl.Videos 1-5**). Regions of interest (ROI) were randomly chosen from 2-4 organoid slices from each line and imaged for 4 minutes at 3.78Hz, a temporal resolution sufficient to detect Ca^++^ events in a time-lapse series (see **Fig. 6E-L** for C5D and **Fig. 6M-T** for C4V). Only ROI with more than 10 neurons were considered for the analysis, as shown in representative maximum projection fluorescence time-lapse image stacks **(Fig. 6C,D)**. The Ca^++^ transients were abolished after tetrodotoxin (TTX) treatment, reflecting the action potential-dependent nature of the recorded neuronal activities (**Video S1-S3**). Spontaneous activity could be detected in cells from both organoids’ conditions, with more active neurons detected in C5D derived organoids compared to C4V (58% ±5.77 vs 39.8% ±4.96 in C4V, **Fig. 6E, K, M, S**).

**Figure 6.**
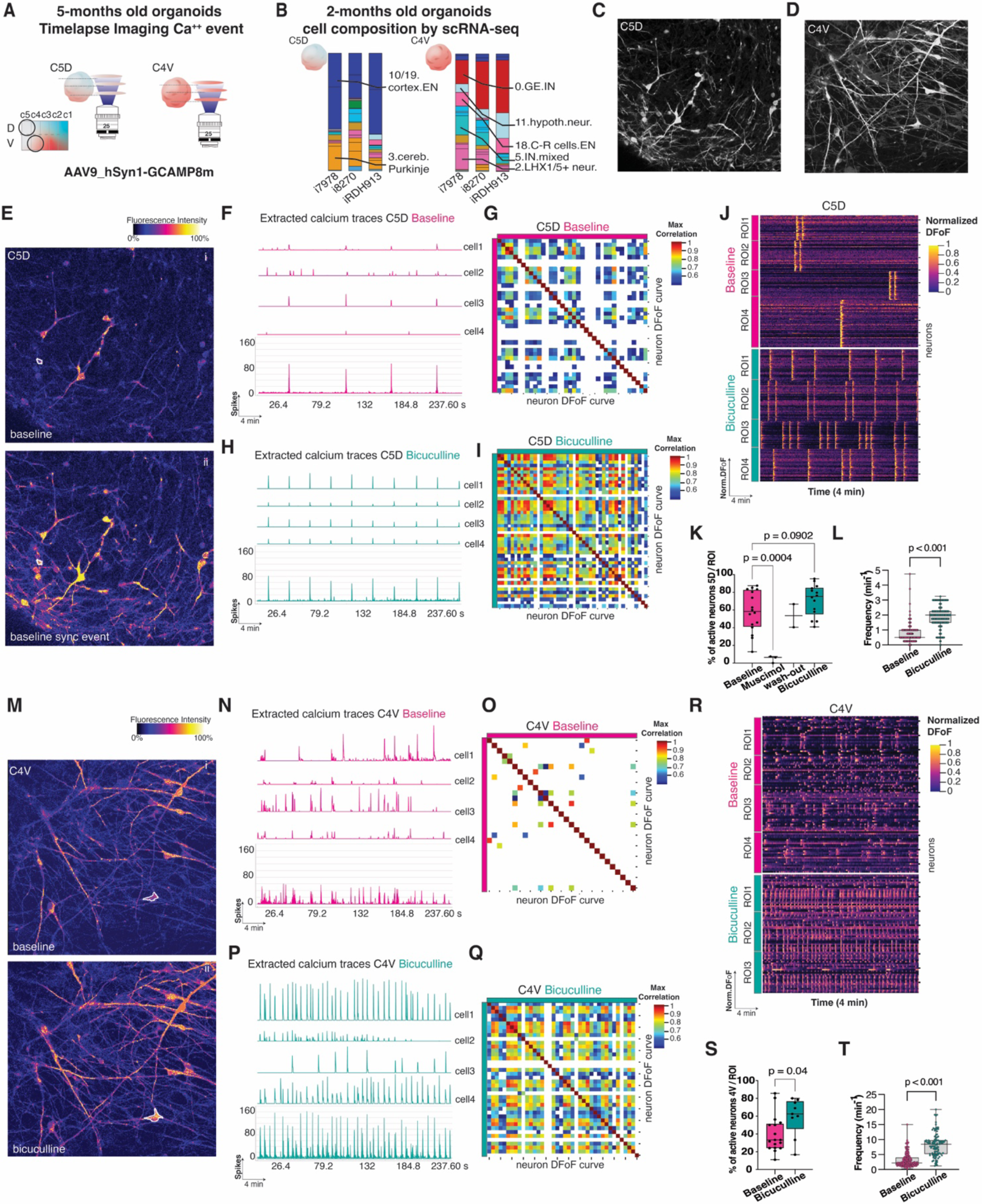
Functional analysis of neuronal signaling dynamics in antero-dorsal (C5D) and antero-ventral organoids (C4V) at 5 months. **A)** Antero-dorsal and antero-ventral organoids were sliced for functional studies with the genetically encoded calcium indicator AAV9-hSYN1-GCamp8m at 5-months in culture. **B)** Cell lineages composition bar plots of 2-months old organoids derived from the 3 different iPSC lines. **C-D)** Maximum fluorescence projection of GCamp8m-labeled neurons in sliced organoids from C5D and C4V (2 weeks after slicing, 4 min time lapse stack). **E)** Changes in GCamp8m fluorescence level in representative single image of C5D organoids at baseline and during a synchronous Ca^++^ transient (sync) (i, ii). **F,H)** Traces of Ca^++^ transients in a C5D organoid at baseline (**F**) and after 50 mM bicuculline exposure (**H**) with aggregated curves (bottom) and 4 representative traces from single neurons (top). **G,I**) Heatmaps of cross-correlations coefficients between normalized curves of each neuron’s activity from a 5D organoids sample in a representative field at baseline (**G**) and after bicuculline (**I**). Note the increase in synchronously active neurons after bicuculline. **J)** Raster plot of normalized calcium traces over time before and after bicuculline treatment in 4 different regions of interest (ROI). **K)** Box plot of percentage of active neurons per ROI per 4 min recording period at baseline, after 25 mM Muscimol and bicuculline treatments in C5D organoids. Significance by unpaired t-test (n=2 iPSC lines, 2-3 ROIs from 3 independent organoids per line). **L)** Box plot of Ca^++^ event frequency in C5D organoids at baseline and after bicuculline treatment (peaks per min, in t = 4 min). All boxplots show median, interquartile box and min/max whiskers. **M)** Single images of GCamp8m fluorescence level in C4V organoids before (i) and after treatment with 50 mM bicuculline (ii). **N,P)** Traces of Ca++ transients in a C4V organoid at baseline (**N**) and after 50 mM bicuculline exposure (**P**) with aggregated curves (bottom) and 4 representative traces from single neurons (top). **O,Q**) Heatmaps of cross-correlations coefficients between normalized curves of each neuron’s activity from a C4V organoids sample in a representative field at baseline (**O**) and after treatment with bicuculline (**Q**). **R)** Raster plot of normalized calcium traces over before and after bicuculline treatment in 4 representative ROIs. **S)** Box plots of percentage of active neurons per ROI at baseline and after bicuculline treatment in C4V organoids. Significance by unpaired t-test (n=3 iPSC lines, 2-3 ROIs from 3 independent organoids per line). **T)** Box plot of Ca++ event frequency across all recorded neurons in C4V derived organoids (peaks per min, in t = 4 min).

A major difference between the 2 conditions was the apparent higher number of active neurons as well as their synchronous activity (as measured by temporal cross-correlation signal analysis of Ca++ transients) in C5D (dorsal cortex) recorded neurons compared to C4V (basal ganglia) neurons at baseline (**Fig. 6F,G vs 6N,O**), suggesting different types of neuronal networks. We hypothesized that this difference could be related to the presence of inhibitory synaptic transmission in 4V-derived organoids, which contain higher proportion of GABAergic cells (GE IN, **Fig.6B**, **Fig.S10A**), and significantly increased expression of the enzymes GAD1/2 that catalyze the production of GABA compared to C5D, along with increased expression of GABA_A_ and Glycine receptor subunits known to be expressed in fetal brain (**Fig. S10B-D**). Thus, we performed pharmacological perturbations to test the importance of the GABAergic system in the neuronal network of each condition. First, C5D-organoids treated with Muscimol, a selective GABAA agonist, showed a major decrease in activity, with only 4.6% of active cells exhibiting Ca^++^ transients compared to an average of ∼55% neurons active in each 4 min recording period at baseline (**Fig. 6K, Video S4-5**). Following a 10 min wash, neuronal activity was restored with an equivalent number of active neurons compared to baseline. This result suggested that, at this developmental stage (5 months of differentiation), the “GABA-shift” has already occurred, and GABA exposure leads to hyperpolarization in recorded neurons ^62^.

This finding suggested but did not demonstrate that high GABA tone in C4V-neurons at baseline was responsible for the reduced neuronal activity and limited synchronicity compared to C5D-neurons. To corroborate this, we disrupted inhibitory transmission by treating both C5D and C4V organoid slices with bicuculline (BIC, 50 µM), a potent GABAA receptor antagonist. A single pulse of BIC-treatment significantly increased the number of active neurons, frequency of calcium transients and synchronicity per field in both conditions (**Fig. 6H-L, P-T**). Synchronous events that were recorded abundantly in C5D derived neurons at baseline (**Fig. 6J**) and were rare in 4V derived neurons at baseline (**Fig. 6R**), became synchronous in both conditions after BIC treatment, as shown by the raster plots (**Fig. 6J,R**) and by the similarity of cross-correlation heat maps of neuronal activity between cortical and basal ganglia organoids after bicuculline treatment (**Fig. 6I, Q**). In summary, at ∼5 months of development, organoids showed spontaneous electrical activity, but ventral telencephalic/basal ganglia organoids displayed only rare synchronous events as compared to dorsal cortex. We showed that this was attributable to an increased GABAergic tone, which agreed with the higher proportion of GABAergic neurons present in this region.

## Discussion

Defining the molecular rules controlling the developmental blueprint of the human brain remains an important endeavor. In this study, we demonstrated how the combinatorial effect of early morphogen gradients activating SHH and WNT signaling in orthogonal directions for 5 days were sufficient to specify the major regional lineages of the fetal telencephalon, diencephalon, midbrain, cerebellum, and pons in organoids from multiple human iPSC lines. We linked this diversity to the activation of different gene programs that responded to defined extracellular concentrations of WNT and SHH. This validated the idea that intersections of continuous morphogen gradients drive lineage specification in the human brain. However, iPSC lines from unrelated individuals were biased in their response to morphogens, revealing an influence of intrinsic factors on brain regional and cellular specification.

Previous studies modeled early rostro-caudal and ventral patterning using a WNT agonist in monolayers of neural progenitors derived from the H9 human embryonic stem cell line in a microfluidic device ^18^, or using natural diffusion of SHH from a localized cellular source into H9-derived forebrain organoids ^17^ or gradient chips ^19,63^. The Duo-MAPS’ innovation relies on the capability of generating dual morphogen gradients in a 3D space by passive diffusion, exposing multiple EBs simultaneously to different morphogen environments where one morphogen influences the cellular response to the other. It would be important to test other morphogen combinations, as well as different concentrations or timing of SHH/WNT exposure which could yield different cellular yield and distribution in patterned organoids. The Duo-MAPS’ device design can be easily adjusted to accommodate diffusion dynamics related to other molecules of different sizes and properties to test other morphogen combinations, environmental exposure and drug screening.

Five days of exposure to gradients of the WNT agonist CHIR and the secreted neuropeptide SHH in the earlier stages of neural induction was sufficient to generate a diversity of glutamatergic and GABAergic cell types that populate a variety of regions in the human brain. The dual gradient triggered the activation of other signaling pathways, like BMP, RA, and FGF, mimicking secondary signaling from the anterior neural ridge, or the isthmic organizer at the midbrain-hindbrain boundary. The interplay of WNT and SHH provided the foundation for a cascade of later events. We deciphered the transcriptional programs induced by these morphogens at various concentration steps, identifying spatial modules and TFs with sustained regulatory action on the specification of cell fate, and confirmed their consistent distribution across diverse regions of the human fetal brain *in vivo*^42^, underscoring the fidelity of our *in vitro* model.

Forebrain lineage required the absence of WNT to be robustly specified, as in lower vertebrates, and thus absence of SHH and WNT was key to the initial development of the cortical plate ^64^. On the opposite pole, the ventral-posterior lineage of dopaminergic neurons depended on dual activation of both WNT and SHH signaling in human neural stem cells ^33,65^. Interestingly, cerebellar fates also showed a unique topological location across all dorsalized positions, mimicking the cerebellar development *in vivo* from the roof plate ^66^. Cerebellar fates were driven in organoids by dorsally expressed TFs such as TFAP2A/TFAP2B/POU4F1/ATOH1, as well as by *PAX6* (expressed in granule neuron precursors)*, OLIG3*, *GLI3,*and *LHX1/5*, all of which are associated with the development of the cerebellum *in vivo* ^66-68^. Some of these genes are activated by WNT such as TFAP2A, TFAP2B, and other repressed or independent from it, such as ATOH1 and PAX6. It would be important to test the influence of other signaling molecules such as FGF8 or RA signaling to further restrict topologically the specification of those fates ^69^.

As iPSC lines and organoids maintain the genetic background of the individual of origin, the system provides the tools for investigating the effect of each individual genetic background upon early lineage specification. We found a surprising degree of variation in the patterning response between lines, with lines exhibiting hypo- and hyper-responsiveness to WNT and SHH. Although more work is needed to understand the causes of this line-to-line variation across experimental batches and clones, the finding implied that brain patterning might be slightly different across individuals, accounting for inter-individual variations in relative size, position, and cell type composition of various brain regions ^70^ with potential implications for risk to various brain disorders. Our results also implied that differential morphogen response could account for a large fraction of the variation observed across lines, individuals, laboratories, and protocols in organoid studies, complementing and extending observations made by other groups ^71^. Using enough iPSC clones, replicates, and conditions will be necessary to test how iPSC phenotypic, transcriptomic, genomic, and epigenetic profiles relate to each iPSC line’s ability to respond to morphogens.

Similarity between broader cell types in organoids and fetal human brain was not limited to the transcriptome but was also reflected in functional activity. The synchronous Ca^++^ events observed in the cortex were indicative of more mature and interconnected neural networks, where neurons dynamically coordinate their firing to process information ^72^. This suggested that our culture system at 5 months allowed the formation of such neural networks. We show that the much lower synchronicity in the basal ganglia was driven by their higher GABAergic tone, reflecting fundamental differences in network organization ^73^. This is consistent with the prevalence of GABAergic neurons in basal ganglia *in vivo* and with prior observations that reducing intrastriatal GABA inhibition synchronizes striatal dynamics, leading to involuntary movements ^74,75^. Our approach provided a platform for studying the molecular mechanisms by which different brain regions develop and connect to each other. Furthermore, common neurodevelopmental disorders, such as autism and schizophrenia, involve more than one brain region and necessitate a platform for modeling disease pathogenesis and screening new potential therapeutic targets across regions.

## Limitation of the study

There were several limitations to our approach. The connections between TFs active at day 9 in response to patterning and those maintaining lineage commitment at day 64 were based upon correlations and will require targeted perturbation experiments for definitive demonstration. Finally, although our results demonstrate an instructive role of morphogens in the specification of lineage diversity in the brain, we observed differences in responsiveness between iPSC lines of different genetic backgrounds. While this is perhaps the most interesting result of the study, it limits the topological accuracy of the approach to generate a specific lineage at a precise position, since each line showed different lineage compositions under equivalent morphogen exposure. Yet, this result underlined that intrinsic cellular differences may shape fundamental fate decisions induced by the environment. The iPSC and organoid models provide the tools to understand the nature of these differences.

## Acknowledgments

We thank the members of the Vaccarino laboratory for discussions and contributions to methods. We thank Livia Tomasini for technical support and Monica Mleczek for administrative support.

We thank Prof. Stefano Vicini for discussions on the neuronal activity data and Michael Strickler for research computing support. We thank G. Wang and C. Castaldi, and the Yale Center for Genome Analysis, for library preparation, deep sequencing and Cell Ranger analysis.

We acknowledge the following grant support: National Institute of Mental Health grant no. RF1 MH 123978 (F.M.V. and A.L.); the Kavli Institute for Neuroscience at Yale (Innovator award to F.M.V. and A.L.); Simons Foundation award no. 975844 (F.M.V.) and the Yale Child Study Center (Trainee Pilot Research award to S>C).

## Author contributions

F.M.V., A.L. conceived the study and provided funding; S.S., T.-Y. K., A.J. designed and optimized experiments; T.-Y. K. constructed the device and characterized morphogens’ diffusion profiles and stability; L.Y. performed simulation of morphogen gradients and discussed calcium imaging method; S.S. and A.J. generated organoids and processed them for qPCR, RNA-seq, scRNA-seq, and ICC; S.S. performed and analyzed calcium imaging experiments; A.J. and F.W. performed RNA-seq and scRNA-seq bioinformatic analyses; A.J. performed secondary analyses; A.A. and V.S. performed genomic analyses of WGS data; G.M.A. performed HPLC monoamine assays; A.N. performed ICC and quantification analysis; J.M. maintained organoids culture and participate to discussion methods; A.J., S.S., T.-Y. K. generated display items and wrote the manuscript; all authors provided edits and comments on the manuscript.

## Supplemental information

**Document S1.** Figures S1–S10

**Table S1**: Excel file containing additional data too large to fit in a PDF, related to Material and Methods, Figure 1 and Figure 4. Legend: T1: donors and iPSC line metadata; T2: d9 bulk RNA-seq metadata; T3 d64 scRNA-seq sample metadata; T4: d64 10X single cell library cellranger QC.

**Table S2**: Excel file containing additional data too large to fit in a PDF, related to Figures 1, S2, 4 and 5. Legend: T1: d9 bulk RNA-seq DEGs between device positions; T2: d9 bulk RNA-seq DEG between device and bath positions; T3 : d9 variance partition results, T4: d9 morphogen-response patterns.

**Table S3**: Excel file containing additional data too large to fit in a PDF, related to Figures 2, S2, 4 and 5. Legend: d64 scRNA-seq cell metadata.

**Table S4**: Excel file containing additional data too large to fit in a PDF, related to Figure 3. Legend: T1: d64 cluster annotation, T2: d64 cell composition.

**Table S5**: Excel file containing additional data too large to fit in a PDF, related to Figure S7, Figure 5. Legend: T1: d64 scRNA-seq cluster markers, T2: d64 DEGs results, T3: d64 neuronal DEGs for Ca++ samples.

**Table S6**: Excel file containing additional data too large to fit in a PDF, related to Figure 5. Legend: T1: d64 SCENIC cluster-specific TF regulators; T2: d64 SCENIC regulons; T3: d9vd64 correlations d9 TF RNA level with d64 cell percentage.

**Video S1**. Spontaneous calcium activity recording of representative C5D-organoid, related to Figure 6A– L.

**Video S2**. Calcium activity recording of representative C5D-organoid treated with muscimol, related to Figure 6A–L.

**Video S3**. Calcium activity recording of representative C5D-organoid treated with bicuculline, related to Figure 6A–L.

**Video S4**. Spontaneous calcium activity recording of representative C4V-organoid, related to Figure 6M– T.

**Video S5**. Calcium activity recording of representative C4V-organoid treated with bicuculline, related to Figures 6M–T.

## Material and methods (short summary; see STAR methods for full description)

### Mesofluidic device fabrication and characterization of diffusion profiles

The mesofluidic device, made of Polydimethylsiloxane (PDMS) with a glass cover bottom, featured five connected culture areas separated by trapezoid pillars. It was loaded with a hydrogel (1% Agarose, 1.5mg/ml rat tail collagen type 1). Two fluorescent molecules, fluorescein sodium salt (376 Da) and Antonia Red-dextran 20 (20 kDa), were used to track morphogen-like gradient dynamics. COMSOL Multiphysics simulated molecular diffusion across the device.

### Neuronal Differentiation of iPSC lines

We used seven previously generated human male control iPS cell lines ^11,35^. Cells were differentiated into neurons using a dual-SMAD inhibition protocol ^35^ with SB431542 and LDN-193189. Organoids were patterned by adding to the mesofluidic device the caudalizing WNT activator CHIR99021 which inhibits GSK3i, and the ventralizing morphogen SHH (See Supplemental Experimental Procedures and **Table S1**). Immunostaining, quantitative RT-PCR were performed as described in Supplemental Experimental Procedures.

### RNA-seq and scRNA-seq analysis

Samples (see **Table S1**) were processed for RNA-seq and scRNA-seq analysis as described in Supplemental Experimental Procedures. Differential expressed genes in bulk RNA were inferred using the edgeR pipeline and an FDR cut-off of 0.05 was used for all the tests. scRNA-seq was analyzed using functions from the Seurat suite, except for GRN construction (SCENIC) and RNA velocity analysis (scvelo).

### Calcium imaging

By utilizing Gamp8m we recorded calcium activity in 5-months old organoids derived from dorsal C5D and ventral C4V positions in 3 different iPSC lines, to investigate the dynamic activity of neuronal networks. Organoids were sliced, loaded with the genetically encoded calcium indicators under the Synapsin I promoter, and confocal microscopy was used to visualize and quantify changes in calcium fluorescence over time. All waveform activity was abolished by TTX treatment. Pharmacological agents such Bicuculline and Muscimol were used to test how the excitability of neuronal networks was modulated by, respectively, blocking or activating GABA receptors.

## Data availability

This study did not generate new unique reagents or DNA constructs.

Primary cell lines and iPSC lines are shared via the Infinity BiologiX LLC repository (https://ibx.bio) or by request to the lead contact, Dr. Flora M. Vaccarino, with a completed materials transfer agreement.

Datasets reported in this study will be available through the NIMH Data Archive (NDA) under collection #C5071, URL: https://nda.nih.gov/edit_collection.html?id=5071. For reasons of subjects’ privacy, as specified in our consent form, NDA only allows for controlled access to genomic sequence data.

Further information and requests for resources and reagents should be directed to and will be fulfilled by the lead contact, Dr. Flora M. Vaccarino.

## Supplementary Information document S1

**<u>Supplementary Figures S1-S10</u>**

**<u>Related to Main fig 1:</u>**

Fig S1: device diffusion analysis

Fig S2: Duo-MAPS vs Batch culture, morphogen concentration effects by qPCR, variance partition

**<u>Related to Main Fig 2:</u>**

Fig S3: scRNA cluster markers and brain projection, voxhunt

Fig S4: brain atlas expression of markers

Fig S5: comparison with GE and cerebellum focused datasets

**<u>Related to Main Fig 3</u>**

Fig S6: cell composition differences and DEG per positions and HPLC

Fig S7: validation by immunostaining

**<u>Related to Main Fig 4</u>**

Fig S8: Morphogen response pattern across brain cells and individuals

**<u>Related to Main Fig 5</u>**

Fig S9: SCENIC-derived regulons

**<u>Related to Main Fig 6</u>**

Fig S10: results supporting Ca++ imaging

**<u>Supplementary Tables (supplied as separate Excel files):</u>**

**Table S1:** Excel file containing additional data too large to fit in a PDF, related to Material and Methods, Figure 1 and Figure 4. Legend: **T1**: donors and iPSC line metadata; **T2**: d9 bulk RNA-seq metadata; **T3** d64 scRNA-seq sample metadata; **T4**: d64 10X single cell library cellranger QC.

**Table S2:** Excel file containing additional data too large to fit in a PDF, related to Figures 1, S2, 4 and 5. Legend: **T1:** d9 bulk RNA-seq DEGs between device positions; **T2**: d9 bulk RNA-seq DEG between device and bath positions; **T3 :** d9 variance partition results, **T4**: d9 morphogen-response patterns.

**Table S3:** Excel file containing additional data too large to fit in a PDF, related to Figures 2, S2, 4 and 5. Legend: d64 scRNA-seq cell metadata.

**Table S4: T1**: d64 cluster annotation, **T2**: d64 cell composition

**Table S5**: Excel file containing additional data too large to fit in a PDF, related to Figure S7, Figure 5. Legend: **T1**: d64 scRNA-seq cluster markers, **T2**: d64 DEGs results, **T3**: d64 neuronal DEGs for Ca++ samples.

**Table S6**: **T1**: Excel file containing additional data too large to fit in a PDF, related to Figure 5. Legend: d64 SCENIC cluster-specific TF regulators; **T2**: d64 SCENIC regulons; **T3**: d9vd64 correlations d9 TF RNA level with d64 cell percentage.

**<u>Supplementary Videos 1-5</u>**: Ca++ imaging (60 fps)

**Video S1.** Spontaneous calcium activity recording of representative C5D-organoid, related to Figure 6A–L.

**Video S2.** Calcium activity recording of representative C5D-organoid treated with muscimol, related to Figure 6A–L.

**Video S3.** Calcium activity recording of representative C5D-organoid treated with bicuculline, related to Figure 6A–L.

**Video S4.** Spontaneous calcium activity recording of representative C4V-organoid, related to Figure 6M–T.

**Video S5.** Calcium activity recording of representative C4V-organoid treated with bicuculline, related to Figures 6M–T.

---

**Figure S1.**
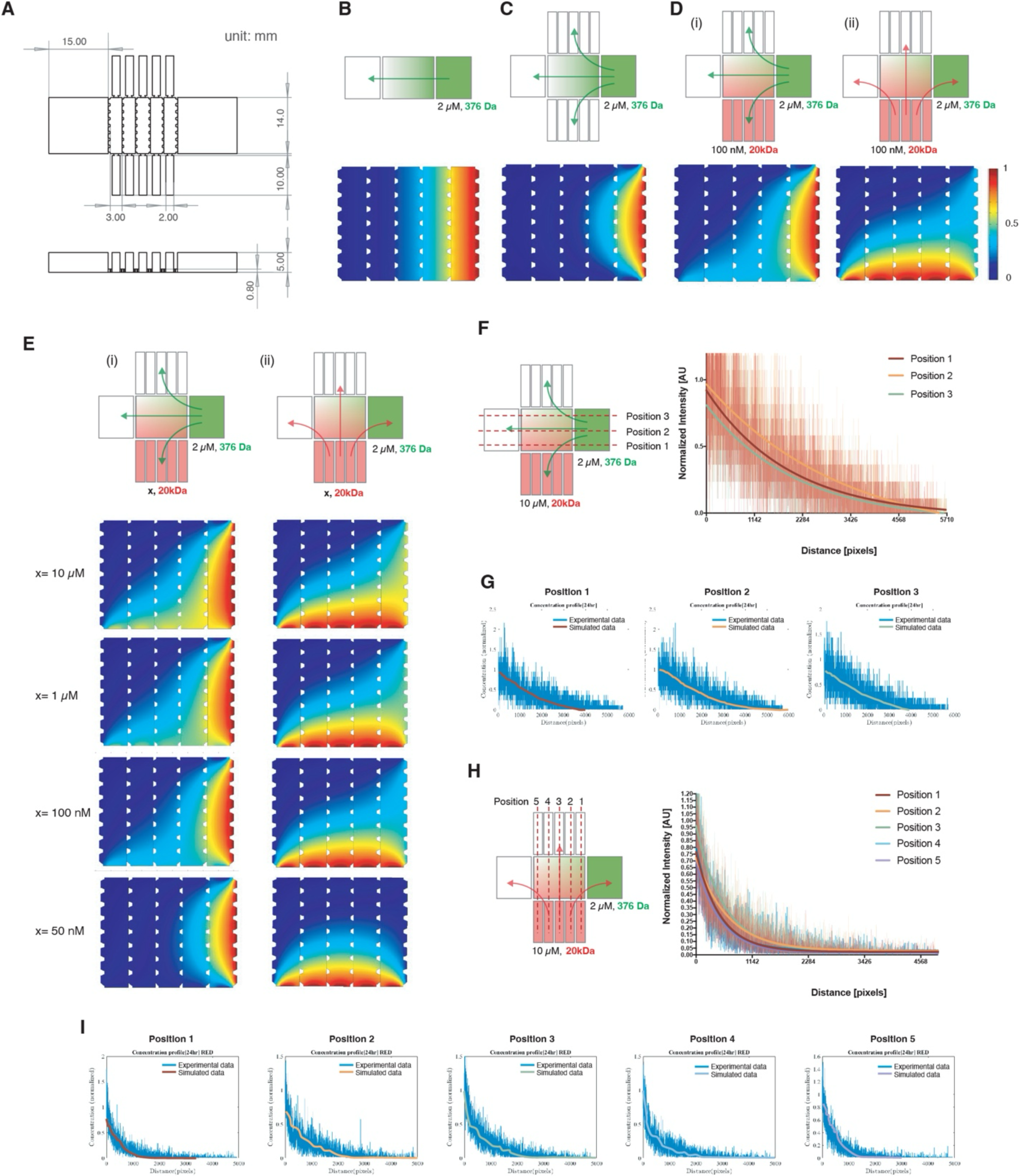
Characterization of morphogens’ diffusion profiles in the Duo-MAPS mesofluidic device. **A)** Schematic and dimensions of the Duo-MAPS. Dimensions of the culture area where EBs are loaded: 15 x 14 x 0.8 mm. **B-D)** Simulated diffusion profiles across the culture area for morphogen of the indicated molecular weights in a unilateral device (**B**) and in the orthogonal device with either one (**C**) or two morphogens (**D**) as shown by top schematics. Diffusion profiles heatmap are given as the relative concentration compared to the source (concentration in the reservoirs). As a result of the presence of orthogonal morphogens and their collision at the corner between the two source reservoirs, the diffusion patterns in both directions were asymmetrically parabolic, slanting towards the side of the other solute’s source reservoir. **E)**. Variations in simulated diffusion profiles in the orthogonal device based on the concentration of the solute. Asymmetrically parabolic diffusion pattern was diminished as the concentration of solutes decreased. **(F)** Experimental and **(G)** simulated diffusion profiles for CHIR along the horizontal axis at three different level (Positions 1, 2, and 3) at 24 hours. **(H)** Experimental and **(I)** simulated diffusion profiles for SHH along the vertical axis at Positions 1, 2, 3, 4, and 5 at 24 hours.

**Figure S2:**
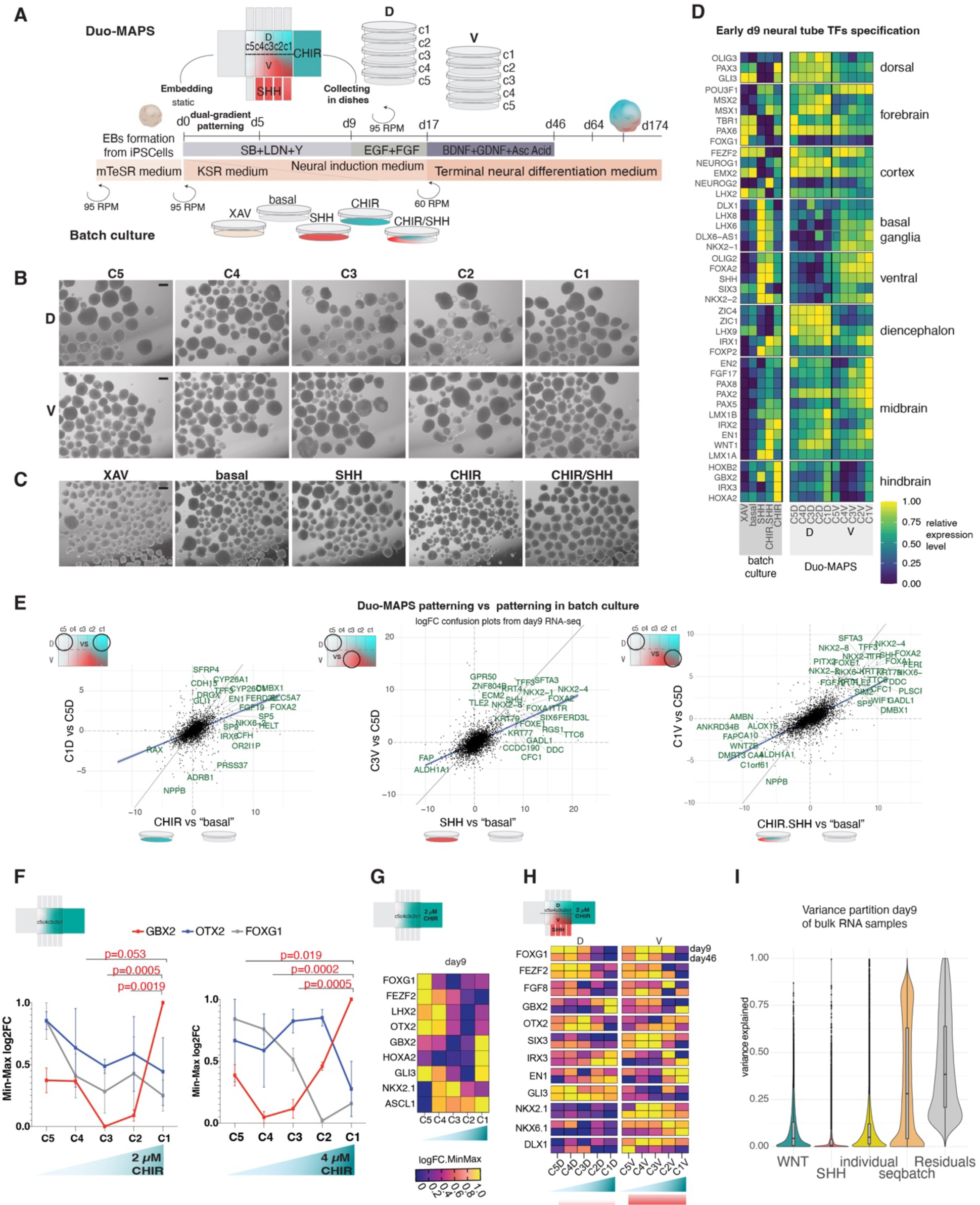
Validation of Duo-MAPS patterning using homogeneous morphogen exposure in rotating dishes. **A)** Detailed study design. In parallel to samples obtained following Duo-MAPS patterning, we applied discrete concentrations of morphogens using addition of morphogens at defined concentration in dishes (’batch method’). For each condition between day 0-5, a single morphogen concentration was tested: XAV (2 µM), basal (no added morphogen, i.e., 0 CHIR + 0 SHH + 0 XAV), SHH (100nM), CHIR.SHH (2 µM CHIR + 100nM SHH), CHIR (2 µM). Concentrations were chosen to reflect extreme exposure conditions in the device, representing either the closest or furthest distances from morphogen sources. See details in **Methods**. **B-C)** Brightfield pictures of different morphogen exposure conditions from the device. (**B**) and from the classical batch culture in dishes at day 46 (**C**). Scale bar, 1mm. **D)** Heatmap of relative expression level at day 9 by bulk RNA-seq for known markers of brain patterning comparing samples exposed to the WNT and SHH gradients in the device (10 positions, C5D-C1V) and samples from the same iPSC lines exposed to single concentrations of the same morphogens added to the culture medium (“batch method”). These concentrations included 2 µM CHIR and 100 nM SHH, separately and in combination, corresponding to the most extreme exposure of the tested morphogens an EB can receive within the device. We also included a “basal” condition with only the dual-SMAD inhibition, and a condition where the WNT signaling pathway was inhibited using the small molecule XAV. All genes passed glm-based ANOVA test across positions with FDR < 0.05 and derived log_2_FC were min-max-transformed (**Methods**). **E)** Confusion plots to evaluate the similarity between differential expression results obtained when comparing 2 device positions (y axis) or when comparing two homogeneous exposures in dishes (x axis). Linear regression lines (blue lines) validated that similar effects of morphogens were obtained in device and classical culture with homogeneous morphogen exposure. This comparison validated that the Duo-MAPS based protocol did not alter the function of morphogens. **F)** RT-qPCR analysis of unilateral gradient of CHIR at day 9 (n=3 iPSC lines) showed how increasing the CHIR99021 concentration at the source reservoir from 2 µM to 4 µM creates a gradient with stronger posteriorizing effect on spheres along the device chambers (C1-C5). The posteriorization effect was evaluated by RT-qPCR of three main TFs defining the antero-posterior axis of the neural tube: GBX2 (posterior), OTX2 (intermediate), and FOXG1 (anterior). Both tested concentrations induced significant increase of GBX2 expression in C1 versus C5, C4, C3 derived samples (Ordinary one-way ANOVA, Tukey’s multiple comparison). An upregulation trend for GBX2 in C2 as well as a downregulation trend of FOXG1 in both C2 and C1, compared to other positions is observable only in 4µM CHIR. **G)** Gene expression profiles of major markers of the antero-posterior and dorsal-ventral axes by RT-qPCR at day 9 (log_2_FC, min-max normalized, n=1 iPSC line) show responsiveness to WNT signaling activation when a *unilateral* gradient of 2 µM CHIR is applied. **H)** Gene expression profiles of major markers of the antero-posterior and dorsal-ventral axes by RT-qPCR following a dual-orthogonal gradient patterning protocol, using 2 µM CHIR and 100 nM SHH at the sources. Results were obtained at day 9 (top row) and day 46 (bottom row) to demonstrate that the regional specification of organoids perdure over time. Comparable expression patterns across device positions at day 9 and day 46 for major brain patterning markers confirmed that 5 days dual-gradient was sufficient to induce stable regional specification (log_2_FC, min-max normalized). **I)** Violin plot showing the distribution of variance explained across all genes for each tested metadata variable (seqbatch: bulk RNA samples were sequenced in two batches, encompassing 2 different groups of individuals, therefore this variable confounds sequencing batch and interindividual variation; individual: remaining variations between individuals).

**Figures S3:**
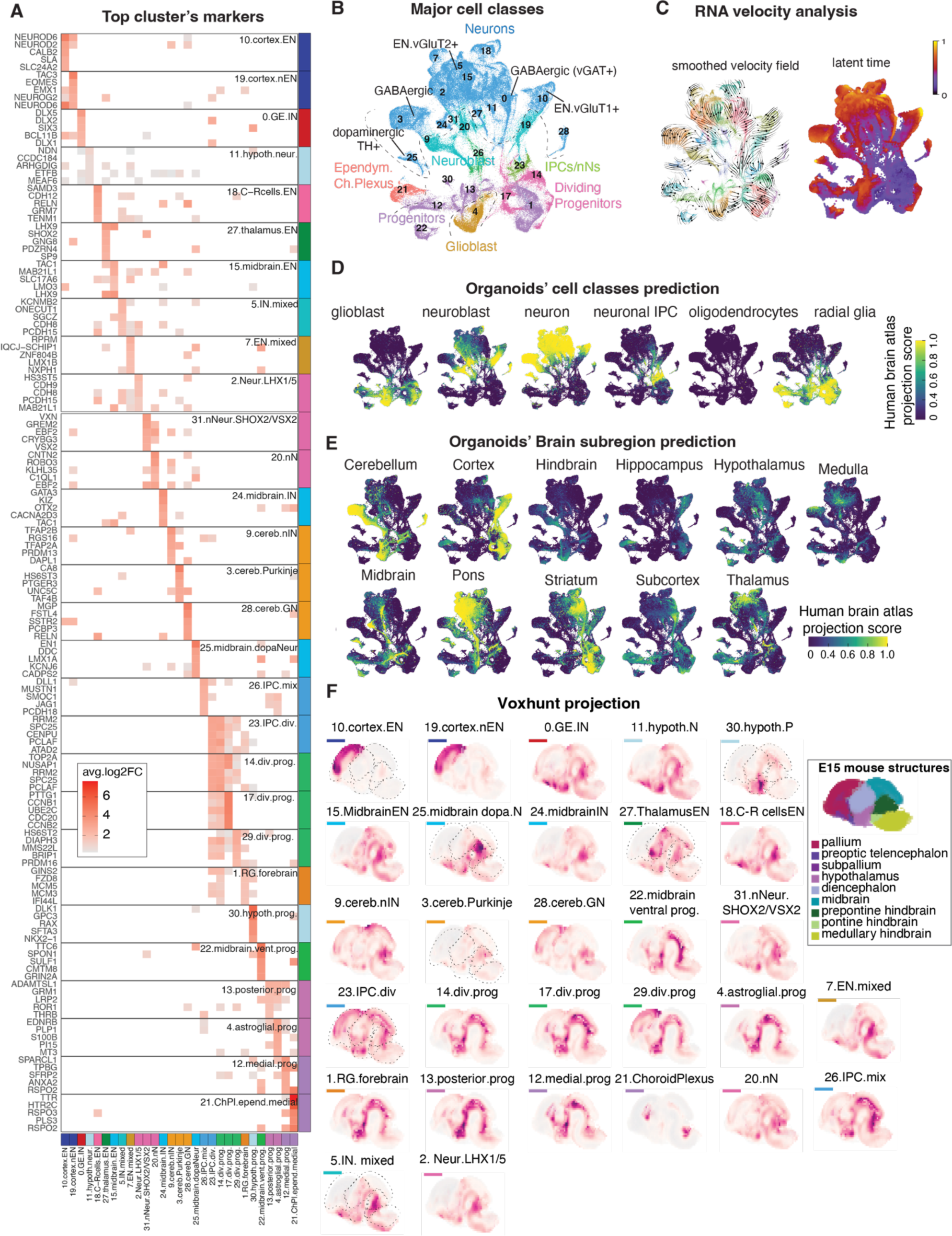
Additional information for scRNA-seq cluster characterization. **A)** Top 5 cluster markers in d64 organoid scRNA-seq data (2-months old). **B)** UMAP plots colored and labelled by major cell classes, neurotransmitter system and regional signature inferred from known cellular marker genes and the Braun et al. fetal brain dataset projection score in B. **C)** Results of RNA velocity analysis using *scvelo* tool showing smoothed velocity field (as vectors showing progression of cell transcription dynamics based on transcript spliced/unspliced ratio per cells) and inferred latent time across all cells on UMAP plots. **D)** UMAP plots of organoid scRNA-seq dataset colored by label transfer score (Seurat, **Methods**) using “cell type” labels from the Braun et al. 2023 ^1^ fetal brain scRNA-seq atlas used as reference. **E)** UMAP plots colored by label transfer score (Seurat, Methods) for “subregion” labels from the Braun et al fetal brain scRNA-seq reprocessed atlas. The “subregion” label in Braun et al. referred to the tissue of origin dissected to obtain the cells. This annotation did not discriminate amongst cell types but showed broad areas of origin. **F)** The Voxhunt tool was used to project organoid clusters’ similarity score on the mouse E15 brain atlas (using the top 20 marker genes per region from mouse atlas, see **Methods**) to give a visual validation of expression similarity across the brain (high=purple, low=grey). The corresponding mouse atlas (bottom) from Voxhunt ^2^ and delineation of regions on some of the plots are included. Organoids clusters are indicated by number, name and lineage color as in panel A.

**Figure Supp 4:**
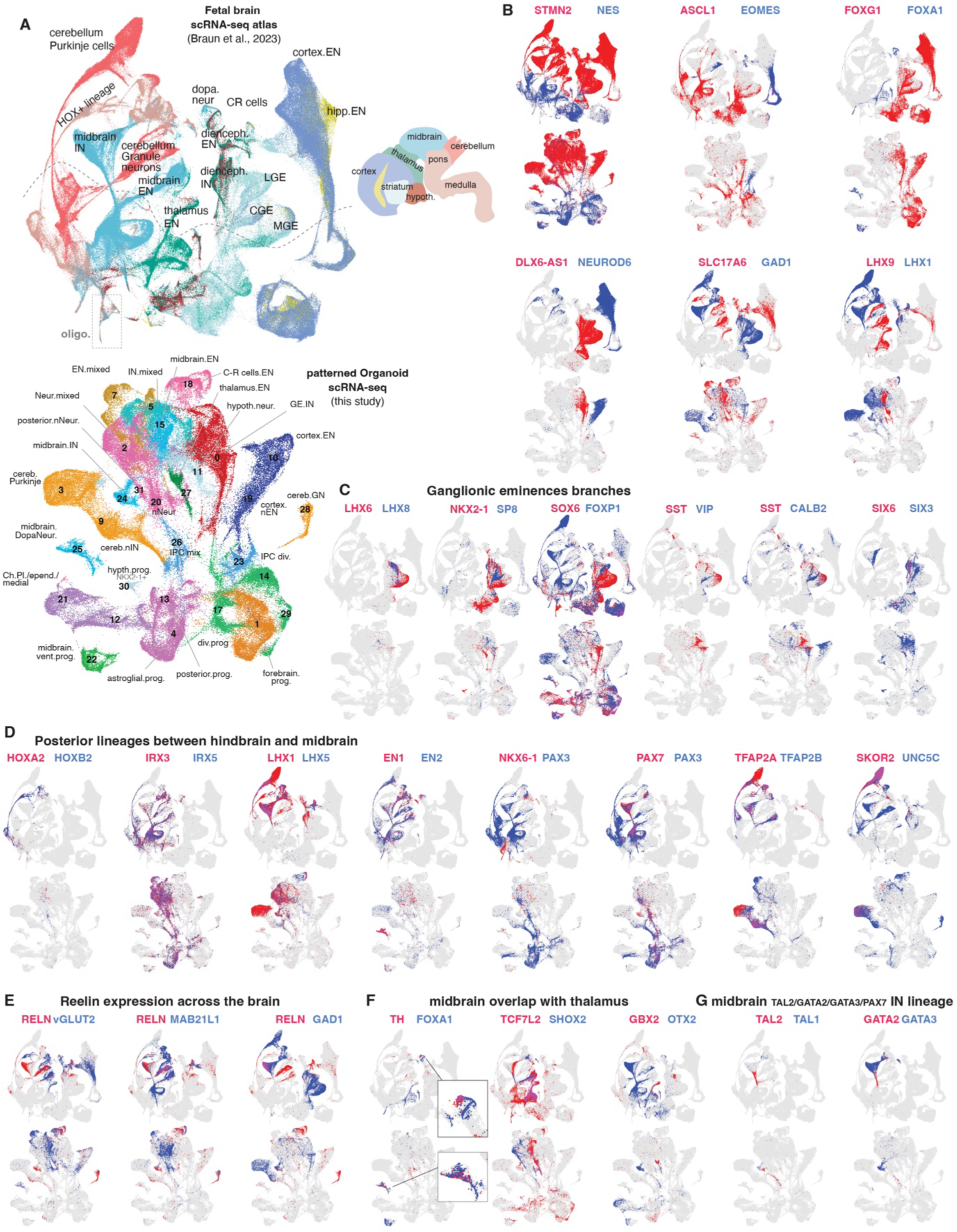
Gene expression profiles marking different regions or lineages in the fetal brain atlas were largely reproduced across cell types generated in patterned organoids. **A** Fetal brain UMAP generated from Braun et al. scRNA-seq dataset, colored by region of origin. Organoid UMAP shown below with cell types annotated as in Fig. 2A. **B-G** Fetal brain UMAPs (top) and organoid UMAP (bottom) showing cells positive for pairs of marker genes (top UMAP, blue vs red, with colocalization in purple). Markers were selected based on their distinctive or overlapping profile across brain regions. While some genes were specific to lineages or subregions (e.g. TAL2/GATA3 in **G**), others were more broad (e.g. posterior regions in **D**). Some genes marked specific lineages in different regions (e.g. FOXP1 in **C** or *FOXA1*, *TCF7L2,* or *GBX2* found in overlap between thalamus and midbrain lineages in **F**). Note how for many gene markers the pattern of co-expression or differential expression across lineages observed in the brain were often reproduced across lineages generated in organoids. The fetal brain atlas brought essential information on marker genes’ exhaustive expression pattern across regions and lineages during early development. While combination of those “markers” could be used to distinguish corresponding lineages generated *in vitro* in scRNA-seq datasets, it should be accompanied by unsupervised annotation methods. *See also **Fig. S3D-F***.

**Figure S5:**
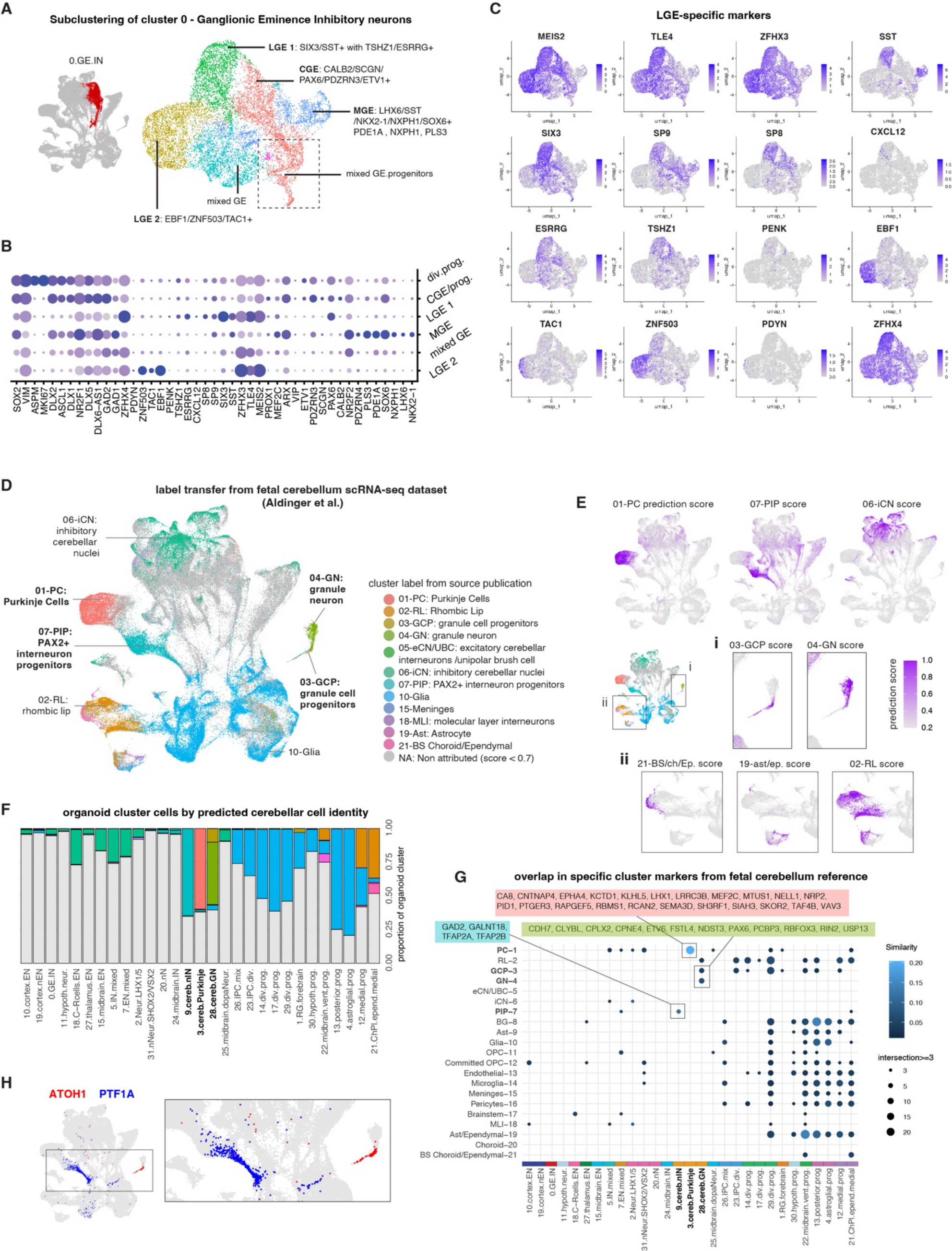
Identification of cell subtypes from ganglionic eminences and cerebellum fates. Related to main Fig. 2. **A)** UMAP plot following subclustering of cells from the ganglionic eminence inhibitory neuron clusters (cluster 0.GE.IN in Fig. 2A) into 6 subclusters, further annotated based on gene expression specific of lateral (LGE), medial (MGE) and caudal (CGE) eminences. **B)** Dotplot of expression level across the subclusters for known LGE-, MGE- and CGE-specific markers. **C)** UMAP plots for gene expression separating different lineages from the LGE related to precursors of medium spiny neurons subtypes (i.e., PDYN+ MSN D1, TSHZ1+ MSN D1 and MSN D2, PENK+) **D)** Organoid cells UMAP (as in Fig. 2A) colored by best matched cell type annotation following label transfer from a human developing cerebellum scRNA-seq dataset from Aldinger et al ^3^. To derive transfer score, the scRNA-seq data of the brain dataset is used as a reference to project the organoid dataset from our study, using transcriptomic similarities between the two datasets. The metadata label from the reference dataset are then transferred to the query based on the labels of its closest neighbors in the reference (see **Methods** and Seurat vignettes). The prediction score (0-1 high similarity) was then used with a threshold of 0.7 to select the best match (cells with score < 0.7 plotted as grey). Cluster legend is from Aldinger et al annotation (color). **E)** Organoid cells UMAP colored by prediction score for the different fetal brain cluster from Aldinger et al. validating the presence of cerebellar-like cells in organoid derived from the device. I) and ii) boxes zoom on specific cell groups as shown on the UMAP. **F)** Barplot showing the proportion of cells in each of the organoid cluster that are labelled by Aldinger et al cell types. Note the specific matching for the 3 clusters in bold. The “Glia” label was attributed mostly with all progenitor cells in organoid (blue). **G)** Dotplot showing the number of cluster-specific marker genes in overlap between the organoid clusters and the Aldinger et al cluster. **H)** Organoid cells UMAP colored by expression for two gene markers of the cerebellar primordium (ATOH1 in the rhombic lip and PTF1A in the ventricular zone).

**Figure S6:**
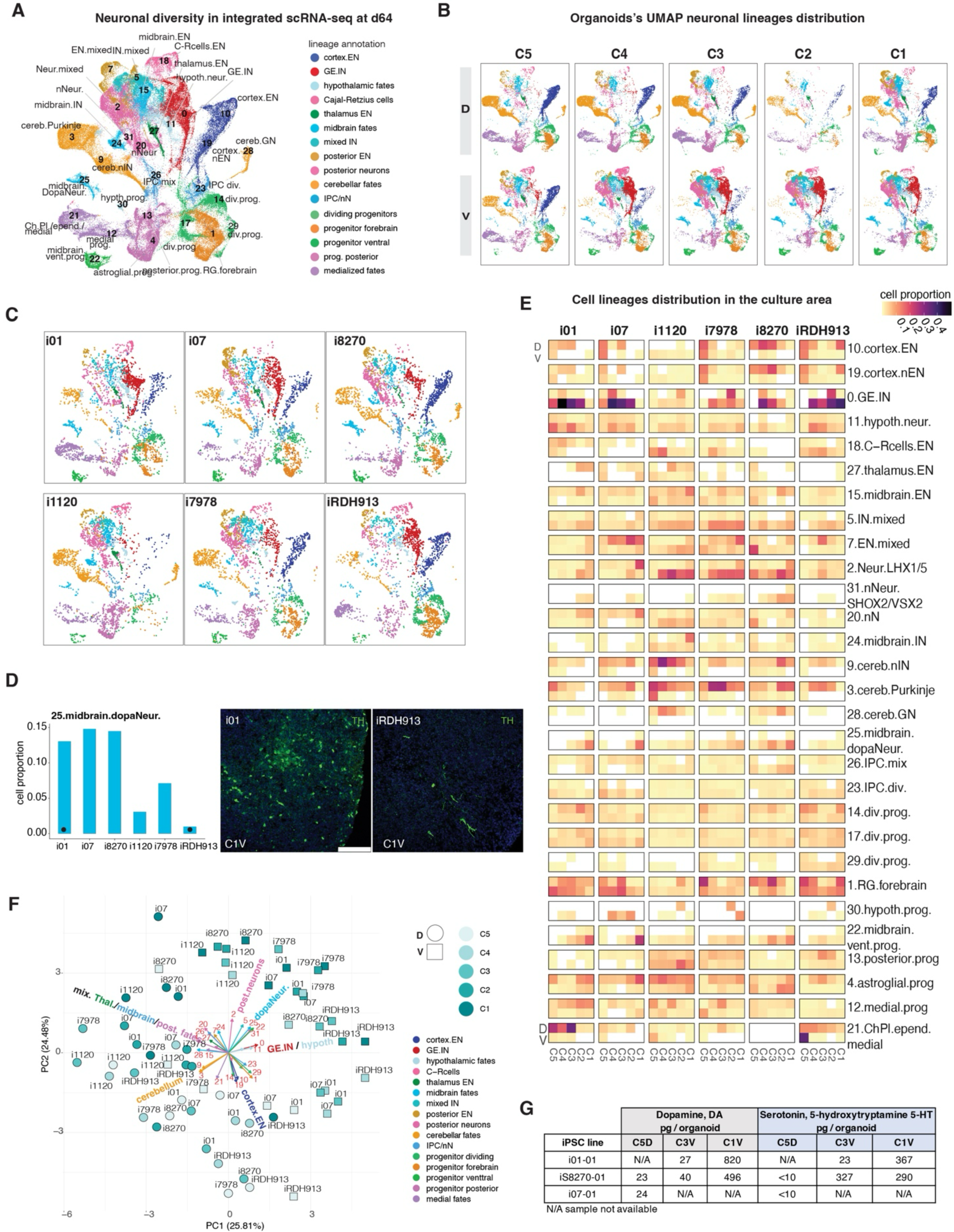
Cell composition analysis for day64 scRNA-seq. **A)** UMAP plot of scRNA-seq on d64 organoids cells colored by main lineage/cell types. **B)** UMAP plots of major identified lineages generated in each device position (average of 6 individuals). Number of cells was subsampled to an equal number by device position (except for C2D which was missing in exp2). **C)** UMAP plots showing the cumulative cell lineages repertoire generated by each iPSC line (6 different individuals= i01, i07, i8270, i1120, i7978, iRDH913) across all device positions. **D)** Percentage of dopaminergic neurons (cluster 25) generated by each iPSC lines in C1V culture position and representative immunofluorescence images showing difference in TH+ dopaminergic cells between two individuals (i01 and RDH913, indicated by a dot) at day64. Scale bar 100 µm **E)** Duo-MAPS device heatmap where D vs V (y axis) and C5 to C1 (x axis) are coordinate representations of the device positions, showing the percentage of cells for each cluster (indicated on the right) across all samples separated by individual lines (e.g., i01, i07, etc.). **F)** Principal component biplot of samples based on cell composition with arrows (colored by lineage) showing contribution to variation of each cluster (cluster number indicated). Dorsal or ventral origin of each sample indicated by different shape, position C5 to C1 indicated by graded color. Cell types generation was primarily influenced by position of origin in the device (see Figure S10 for detailed effect of device position on cell composition) but also dependent on the genetic background of individual lines. **G)** Levels of the neurotransmitters dopamine and serotonin measured by HPLC-EC. C1V-, C3V-, C5D-derived organoids from at least 2 iPSC lines were analyzed (i01, i07, i8270) at day78.**Figure**

**Figure S7.**
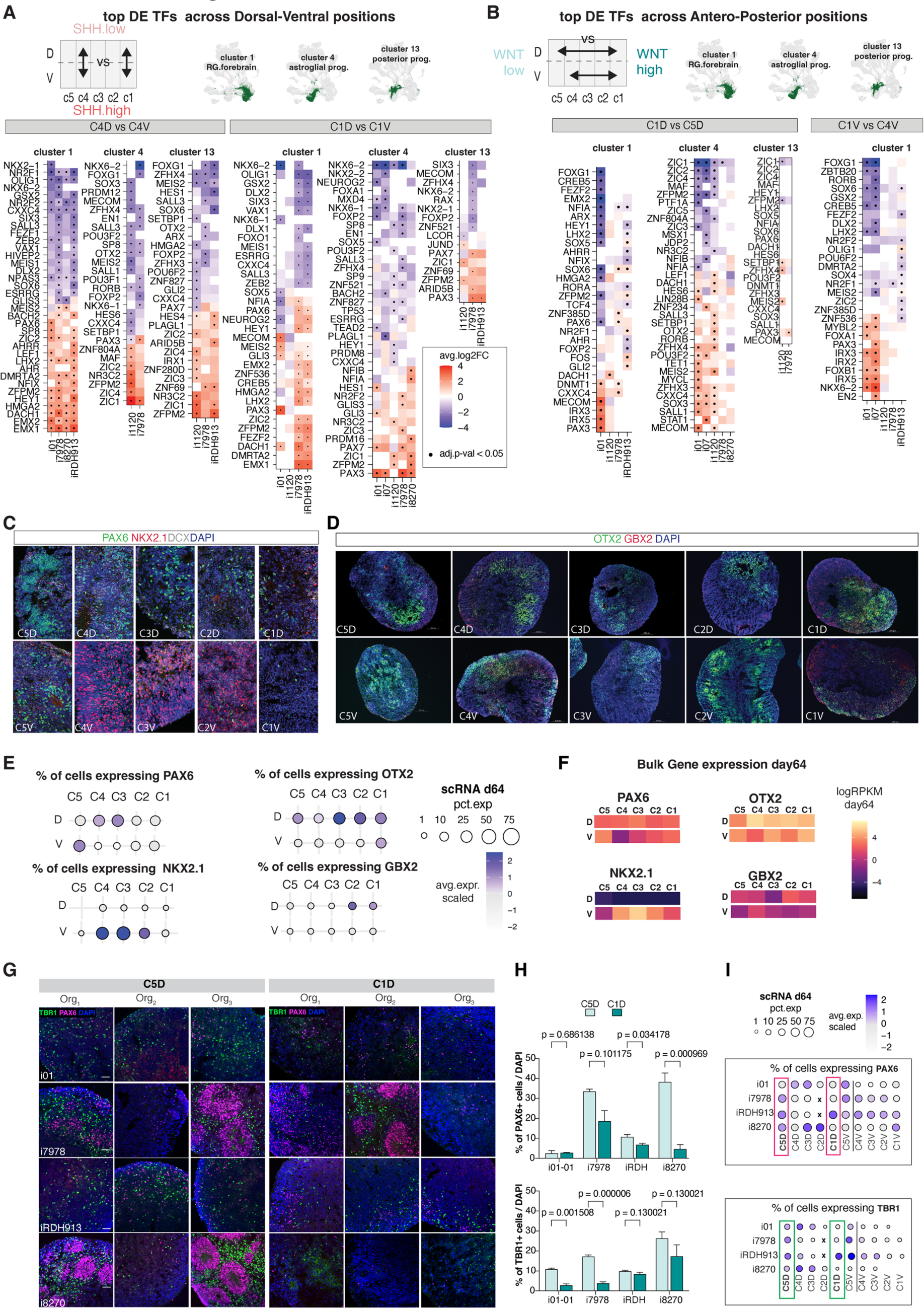
Validation of ventralization or posteriorization of samples using progenitor cells’s differential gene expression and protein marker distribution. **A)** Heatmap of top TFs differentially expressed (DE TFs) between progenitors from SHH^high^ samples and progenitors from SHH^low^ samples (i.e., D vs V positions) using d64 scRNA-seq data separated by chambers, cell types and iPSC lines (average log 2 fold-change (avg.log2FC) showed by color and significance shown by dots for adjusted p-value < 0.05 using Wilcoxon’s rank sum test). To test ventralization at different device positions, differential expression was tested both at the C4 level (left panels) and C1 level (right panels) as shown by top schematic. To test if ventralization was similar across progenitors matching different anteroposterior regional signature, this differential expression was tested separately in 3 progenitor clusters: forebrain RG (cluster 1, exhibiting a mix of cortical and striatal signatures as shown in **main** Fig. 2F) and posterior RGs (clusters 4 & 13, exhibiting a mix of midbrain, cerebellum or pons signature as shown in **main** Fig.2F) with each cluster location indicated in green in the top UMAPs. Finally, this differential expression was tested separately by iPSC line (x axis, note that some lines didn’t present enough cells to be tested, **Methods**). Overall, DE TFs provided strong evidence for progenitors’ long-lasting ventralization following high SHH exposure, marked by strong upregulation of ventral TFs in both C4V and C1V, confirming analysis of cell counts presented in **main** Fig. 3 & **Fig. S6**. This ventralization often involved different sets of TFs in anterior or posterior clusters: anterior progenitor cells ventralized by repressing *LHX2, EMX1/2* and activating *GSX2/DLXs* while posterior progenitor cells ventralized through *PAX7/ZIC1/ZIC4* in posterior ones. **B)** Comparison between WNT high and WNT low positions (i.e., C1 vs C4/5) in either the D portion or V portion in **B.** Changes in expression program in these progenitor clusters between WNT conditions (C1 vs C5) was present but less pronounced. Each DEG test was computed independently in each cell line (Wilcoxon’s rank sum test adjusted p-value < 0.05 indicated by dot; n=30 cells minimum per sample; for genes expressed in at least 10% of cells in either of the samples compared) full result in **Table S5,T2** **C)** Representative immunofluorescence for TFs regional identifier PAX6 (dorsal forebrain), NKX2.1 (GE and hypothalamus), **D)** OTX2 (forebrain-midbrain) and GBX2 (hindbrain) along the device positions confirmed what observed at the gene expression level by both scRNA-seq (**E**) and bulk gene expression (**F**). **G.** Immunofluorescent images comparing organoid sections from C5D and C1D at d64(**A**) (n=4 iPSC lines) 3 different organoids each. Sections were stained with TBR1 (green) and PAX6 (red). **H.** Immunocytochemical quantification of PAX6 (top) and TBR1 (bottom) in 3-4 organoids per individual (BioVoxxel tool, Fiji). P-values by unpaired two-tails t-test. **I.** Corresponding gene expression pattern in scRNA-seq summarized as a pseudobulk (pct.exp= percent of cells expressing the gene, avg.exp.scaled= scaled expression across sample).

**Figure S8:**
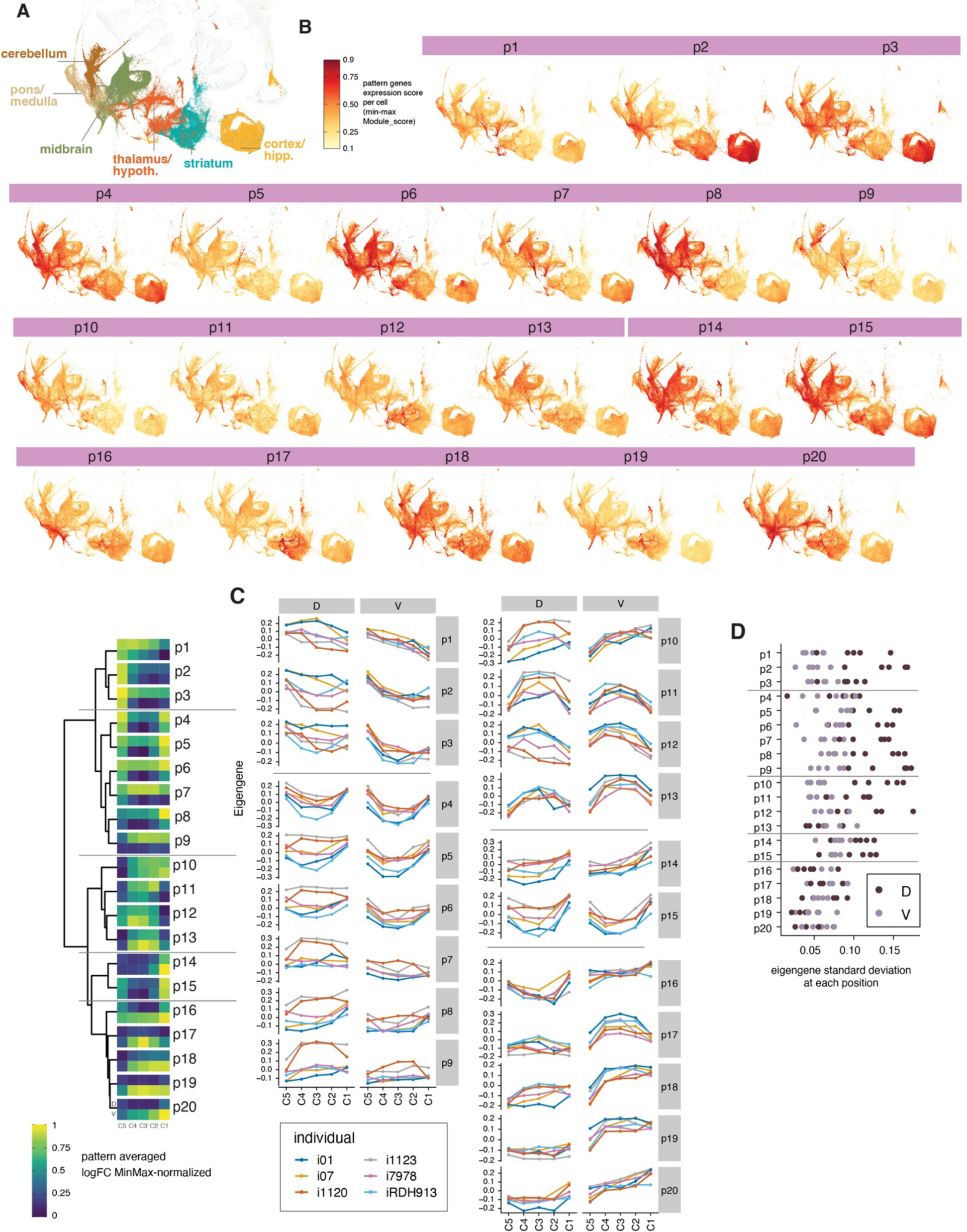
Morphogen response patterns across brain region and individual lines. **A)** UMAP of progenitor cells of the fetal brain atlas colored by region of origin. Progenitor cells = cells defined as radial glia and glial progenitors in Braun et al 2023. **B)** UMAP plots showing the average expression score across single progenitor cells in the fetal brain atlas for each of the 20 morphogen-response spatial modules derived from the device d9 RNA-seq (p1-p20 grouped as in **Main** Fig. 4A, see also in C). Module showed different expression patterns in fetal brain progenitors and across regions. Expression score was computed using Seurat *ModuleScore* function using all genes included in each module (score was min-max normalized across all cells, **Methods**). **C)** DeviceMap showing eigengene expression in each iPSC line for the 20 patterns defined in **Main** Fig. 4A (right side, min-max log2FC average pattern). **D)** Standard deviation of pattern’s eigengene values across the 6 iPSC lines at each device position at day 9 revealed an increased interindividual differences in morphogen-response in the D samples of the device (C5D-C1D). (Note that line i1123 was not analyzed by scRNA-seq at day 64 but is included here at day9.)

**Figure S9:**
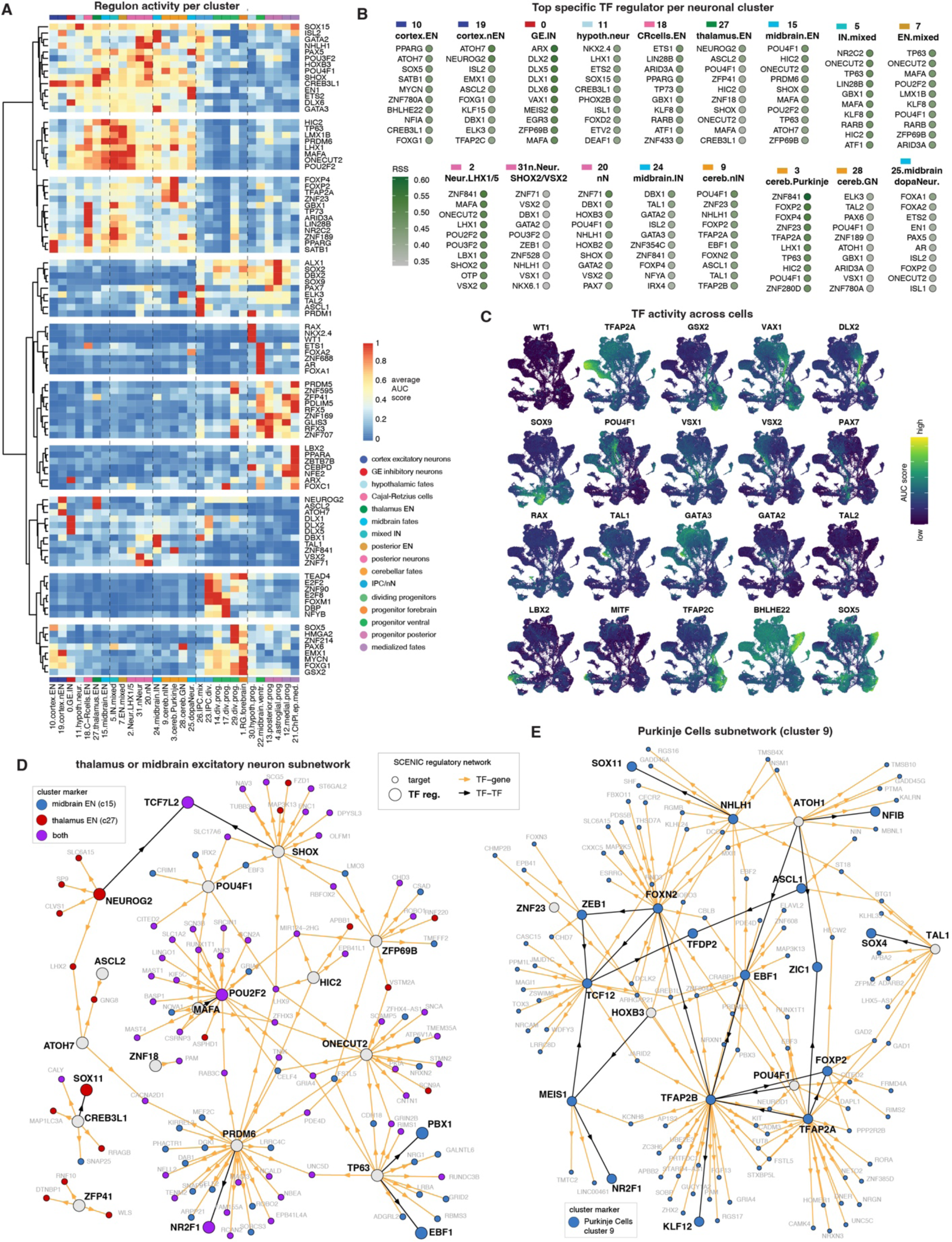
SCENIC TF regulon analysis of d64 scRNA-seq. Clustering of TF regulators identified by SCENIC analysis based on TF’s regulon activity at day 64 (average AUC score per cluster, with AUC score being equivalent to an enrichment of TF’s targets in each cell’s transcriptome, displayed in **C**, see **Methods** ^4^. Heatmap shows the average AUC score per cluster per TF, identifying potential regulators of cell type specific expression. Top 5 cluster-specific TFs based on regulon specificity score (RSS, Methods) for any cluster were plotted. **A)** Top 10 cluster-specific TFs ranked by regulon specificity score (RSS, see **Methods**). See the cluster-specific TFs regulators in **Table S6**. **B)** UMAP plots showing AUC score across cells for selected cluster-specific TFs. **D-E)** SCENIC-derived subnetworks connecting the regulons of the top 10 cluster-specific TFs for vGluT1-positive TCF7L2-positive excitatory neurons (merging 15.midbrain.EN and 27.thalamus.EN, in **D**) and for Purkinje cells (9.cereb.Purkinje in **E**). Only target genes identified as cluster markers of the selected clusters were included (See cluster markers list in **Table S4**). Panel **D** distinguishes cluster markers specific for midbrain (in blue), specific for thalamus EN (in red), or shared by both clusters (in purple). Panel **E** shows Purkinje cells’ cluster markers in blue.

**Figure S10.**
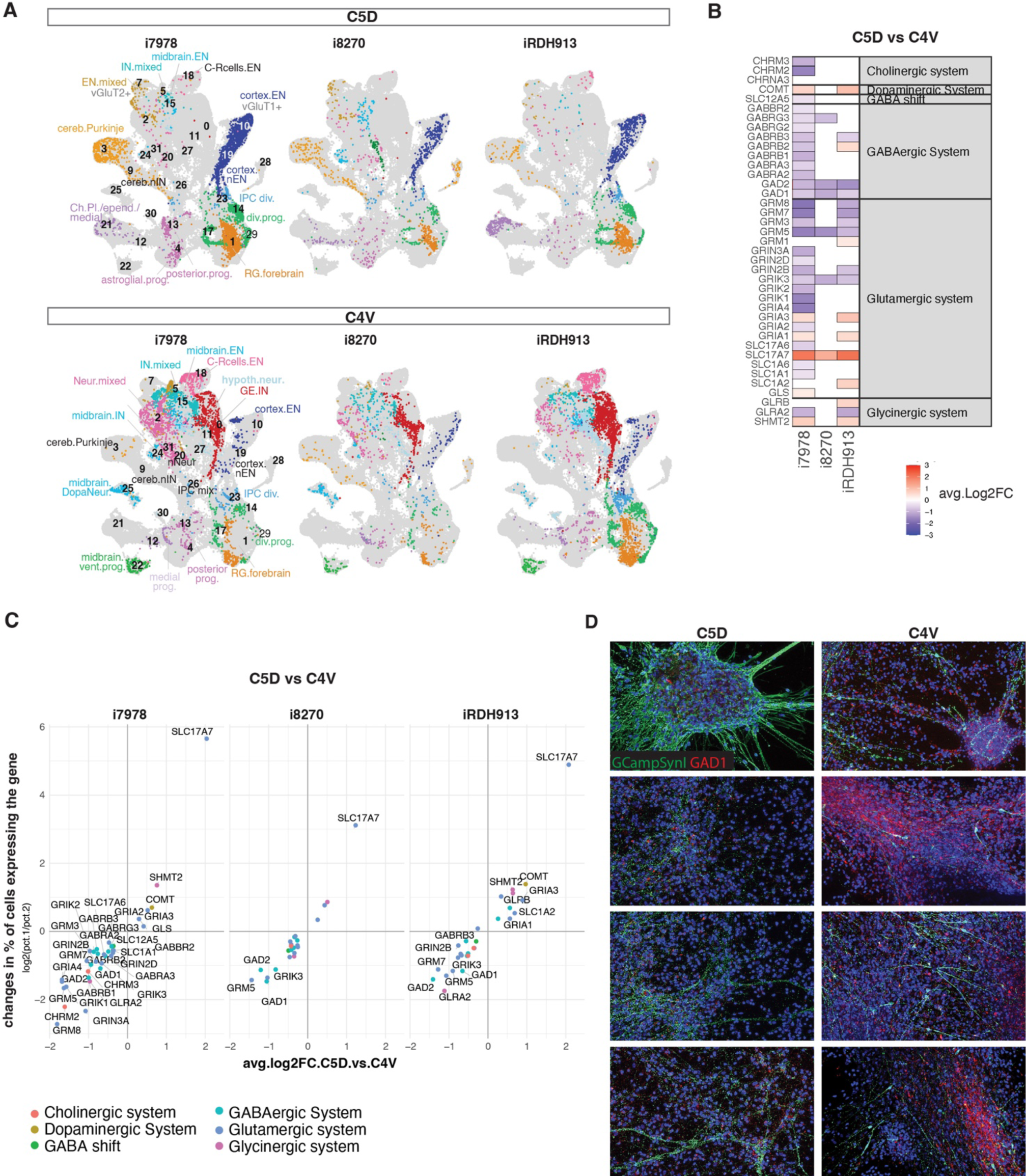
Neuronal composition of samples used for Ca^++^ imaging. Related to Figure 6. **A)** UMAP of cells from d64 scRNA-seq data for the 6 samples analyzed by calcium imaging. **B)** Heatmap of differential expression between C5D and C4V position for each line (x axis) for genes related to neurotransmitter system (y axis). Differential expression was assessed using pseudobulk values across positions. **C)** Dot plots showing the differences in average log2-fold change expression (x axis) and percentage of expression (y axis) for genes in neurotransmitter systems (dot color with genes meeting adjusted p-value < 0.001 named, Wilcoxon rank sum test). **D)** Immunostaining for GAD1 (mouse anti-GAD67 antibody, red) post hoc Ca++ recordings in C5D and C4V sliced organoids, line iRDH913. Each panel represent a different region of interest (ROI), with some showing increased presence of GAD1+cells (red) in C4V position compared to C5D. GCamp8mSynI signal was enhanced with chicken anti-GFP antibody (green); nuclei (blue) are counterstain with DAPI.

## References

1. Ragsdale, C.W., and Grove, E.A. (2001). Patterning the mammalian cerebral cortex. Current opinion in neurobiology 11, 50–58.

2. Wilson, S.W., and Rubenstein, J.L. (2000). Induction and dorsoventral patterning of the telencephalon. Neuron 28, 641–651.

3. Shimamura, K., and Rubenstein, J.L. (1997). Inductive interactions direct early regionalization of the mouse forebrain. Development 124, 2709–2718.

4. Jessel, T.M., and Lumsden, A. (1997). Inductive signals and the assignment of cell fate in the spinal cord and hindbrain. In Molecular and cellular approaches to neural development, W.M. Cowan, T.M. Jessel, and S.L. Zipursky, eds. (Oxford University Press), pp. 290–333.

5. Kudoh, T., Wilson, S.W., and Dawid, I.B. (2002). Distinct roles for Fgf, Wnt and retinoic acid in posteriorizing the neural ectoderm. Development 129, 4335–4346. 10.1242/dev.129.18.4335.

6. Roelink, H., Porter, J.A., Chiang, C., Tanabe, Y., Chang, D.T., Beachy, P.A., and Jessell, T.M. (1995). Floor plate and motor neuron induction by different concentrations of the amino-terminal cleavage product of sonic hedgehog autoproteolysis. Cell 81, 445–455. 10.1016/0092-8674(95)90397-6.

7. Chamberlain, C.E., Jeong, J., Guo, C., Allen, B.L., and McMahon, A.P. (2008). Notochord-derived Shh concentrates in close association with the apically positioned basal body in neural target cells and forms a dynamic gradient during neural patterning. Development 135, 1097–1106. 10.1242/dev.013086.

8. Joksimovic, M., Anderegg, A., Roy, A., Campochiaro, L., Yun, B., Kittappa, R., McKay, R., and Awatramani, R. (2009). Spatiotemporally separable Shh domains in the midbrain define distinct dopaminergic progenitor pools. Proc Natl Acad Sci U S A 106, 19185–19190. 10.1073/pnas.0904285106.

9. Mariani, J., Simonini, M.V., Palejev, D., Tomasini, L., Coppola, G., Szekely, A.M., Horvath, T.L., and Vaccarino, F.M. (2012). Modeling human cortical development in vitro using induced pluripotent stem cells. Proc Natl Acad Sci U S A 109, 12770–12775. 10.1073/pnas.1202944109.

10. Camp, J.G., Badsha, F., Florio, M., Kanton, S., Gerber, T., Wilsch-Brauninger, M., Lewitus, E., Sykes, A., Hevers, W., Lancaster, M., et al. (2015). Human cerebral organoids recapitulate gene expression programs of fetal neocortex development. Proc Natl Acad Sci U S A 112, 15672–15677. 10.1073/pnas.1520760112.

11. Mariani, J., Coppola, G., Zhang, P., Abyzov, A., Provini, L., Tomasini, L., Amenduni, M., Szekely, A., Palejev, D., Wilson, M., et al. (2015). FOXG1-Dependent Dysregulation of GABA/Glutamate Neuron Differentiation in Autism Spectrum Disorders. Cell 162, 375–390. 10.1016/j.cell.2015.06.034.

12. Sakaguchi, H., Kadoshima, T., Soen, M., Narii, N., Ishida, Y., Ohgushi, M., Takahashi, J., Eiraku, M., and Sasai, Y. (2015). Generation of functional hippocampal neurons from self-organizing human embryonic stem cell-derived dorsomedial telencephalic tissue. Nature communications 6, 8896. 10.1038/ncomms9896.

13. Muguruma, K., Nishiyama, A., Kawakami, H., Hashimoto, K., and Sasai, Y. (2015). Self-organization of polarized cerebellar tissue in 3D culture of human pluripotent stem cells. Cell reports 10, 537–550. 10.1016/j.celrep.2014.12.051.

14. Velasco, S., Kedaigle, A.J., Simmons, S.K., Nash, A., Rocha, M., Quadrato, G., Paulsen, B., Nguyen, L., Adiconis, X., Regev, A., et al. (2019). Individual brain organoids reproducibly form cell diversity of the human cerebral cortex. Nature 570, 523–527. 10.1038/s41586-019-1289-x.

15. Xiang, Y., Tanaka, Y., Cakir, B., Patterson, B., Kim, K.Y., Sun, P., Kang, Y.J., Zhong, M., Liu, X., Patra, P., et al. (2019). hESC-Derived Thalamic Organoids Form Reciprocal Projections When Fused with Cortical Organoids. Cell stem cell 24, 487–497 e487. 10.1016/j.stem.2018.12.015.

16. Gabriel, E., Albanna, W., Pasquini, G., Ramani, A., Josipovic, N., Mariappan, A., Schinzel, F., Karch, C.M., Bao, G., Gottardo, M., et al. (2021). Human brain organoids assemble functionally integrated bilateral optic vesicles. Cell stem cell 28, 1740–1757 e1748. 10.1016/j.stem.2021.07.010.

17. Cederquist, G.Y., Asciolla, J.J., Tchieu, J., Walsh, R.M., Cornacchia, D., Resh, M.D., and Studer, L. (2019). Specification of positional identity in forebrain organoids. Nat Biotechnol 37, 436–444. 10.1038/s41587-019-0085-3.

18. Rifes, P., Isaksson, M., Rathore, G.S., Aldrin-Kirk, P., Moller, O.K., Barzaghi, G., Lee, J., Egerod, K.L., Rausch, D.M., Parmar, M., et al. (2020). Modeling neural tube development by differentiation of human embryonic stem cells in a microfluidic WNT gradient. Nat Biotechnol 38, 1265–1273. 10.1038/s41587-020-0525-0.

19. Xue, X., Kim, Y.S., Ponce-Arias, A.I., O’Laughlin, R., Yan, R.Z., Kobayashi, N., Tshuva, R.Y., Tsai, Y.H., Sun, S., Zheng, Y., et al. (2024). A patterned human neural tube model using microfluidic gradients. Nature 628, 391–399. 10.1038/s41586-024-07204-7.

20. Zagorski, M., Tabata, Y., Brandenberg, N., Lutolf, M.P., Tkacik, G., Bollenbach, T., Briscoe, J., and Kicheva, A. (2017). Decoding of position in the developing neural tube from antiparallel morphogen gradients. Science 356, 1379–1383. 10.1126/science.aam5887.

21. Scuderi, S., Altobelli, G.G., Cimini, V., Coppola, G., and Vaccarino, F.M. (2021). Cell-to-Cell Adhesion and Neurogenesis in Human Cortical Development: A Study Comparing 2D Monolayers with 3D Organoid Cultures. Stem cell reports 16, 264–280. 10.1016/j.stemcr.2020.12.019.

22. Shimamura, K., Hartigan, D.J., Martinez, S., Puelles, L., and Rubenstein, J.L.R. (1995). Longitudinal organization of the anterior neural plate and neural tube. Development 121, 3923–3933.

23. Rallu, M., Corbin, J.G., and Fishell, G. (2002). Parsing the prosencephalon. Nat Rev Neurosci 3, 943–951.

24. Rallu, M., Machold, R., Gaiano, N., Corbin, J.G., McMahon, A.P., and Fishell, G. (2002). Dorsoventral patterning is established in the telencephalon of mutants lacking both Gli3 and Hedgehog signaling. Development 129, 4963–4974.

25. Griffiths, S.C., Schwab, R.A., El Omari, K., Bishop, B., Iverson, E.J., Malinauskas, T., Dubey, R., Qian, M., Covey, D.F., Gilbert, R.J.C., et al. (2021). Hedgehog-Interacting Protein is a multimodal antagonist of Hedgehog signalling. Nature communications 12, 7171. 10.1038/s41467-021-27475-2.

26. Allen, B.L., Song, J.Y., Izzi, L., Althaus, I.W., Kang, J.S., Charron, F., Krauss, R.S., and McMahon, A.P. (2011). Overlapping roles and collective requirement for the coreceptors GAS1, CDO, and BOC in SHH pathway function. Dev Cell 20, 775–787. 10.1016/j.devcel.2011.04.018.

27. Shi, X., Zhang, Z., Zhan, X., Cao, M., Satoh, T., Akira, S., Shpargel, K., Magnuson, T., Li, Q., Wang, R., et al. (2014). An epigenetic switch induced by Shh signalling regulates gene activation during development and medulloblastoma growth. Nature communications 5, 5425. 10.1038/ncomms6425.

28. Abu-Khalil, A., Fu, L., Grove, E.A., Zecevic, N., and Geschwind, D.H. (2004). Wnt genes define distinct boundaries in the developing human brain: implications for human forebrain patterning. J Comp Neurol 474, 276–288. 10.1002/cne.20112.

29. Assimacopoulos, S., Grove, E.A., and Ragsdale, C.W. (2003). Identification of a Pax6-dependent epidermal growth factor family signaling source at the lateral edge of the embryonic cerebral cortex. J Neurosci 23, 6399–6403.

30. Harrison-Uy, S.J., and Pleasure, S.J. (2012). Wnt signaling and forebrain development. Cold Spring Harbor perspectives in biology 4, a008094. 10.1101/cshperspect.a008094.

31. Martynoga, B., Morrison, H., Price, D.J., and Mason, J.O. (2005). Foxg1 is required for specification of ventral telencephalon and region-specific regulation of dorsal telencephalic precursor proliferation and apoptosis. Dev Biol 283, 113–127. 10.1016/j.ydbio.2005.04.005.

32. Hu, D., Young, N.M., Li, X., Xu, Y., Hallgrimsson, B., and Marcucio, R.S. (2015). A dynamic Shh expression pattern, regulated by SHH and BMP signaling, coordinates fusion of primordia in the amniote face. Development 142, 567–574. 10.1242/dev.114835.

33. Ye, W., Shimamura, K., Rubenstein, J.L., Hynes, M.A., and Rosenthal, A. (1998). FGF and Shh signals control dopaminergic and serotonergic cell fate in the anterior neural plate. Cell 93, 755–766. 10.1016/s0092-8674(00)81437-3.

34. O’Leary, D.D., Chou, S.J., and Sahara, S. (2007). Area patterning of the mammalian cortex. Neuron 56, 252–269. S0896-6273(07)00772-6 [pii]10.1016/j.neuron.2007.10.010.

35. Jourdon, A., Wu, F., Mariani, J., Capauto, D., Norton, S., Tomasini, L., Amiri, A., Suvakov, M., Schreiner, J.D., Jang, Y., et al. (2023). Modeling idiopathic autism in forebrain organoids reveals an imbalance of excitatory cortical neuron subtypes during early neurogenesis. Nat Neurosci 26, 1505–1515. 10.1038/s41593-023-01399-0.

36. Bouwmeester, T., Kim, S., Sasai, Y., Lu, B., and De Robertis, E.M. (1996). Cerberus is a head-inducing secreted factor expressed in the anterior endoderm of Spemann’s organizer. Nature 382, 595–601. 10.1038/382595a0.

37. Houart, C., Caneparo, L., Heisenberg, C., Barth, K., Take-Uchi, M., and Wilson, S. (2002). Establishment of the telencephalon during gastrulation by local antagonism of Wnt signaling. Neuron 35, 255–265. 10.1016/s0896-6273(02)00751-1.

38. Hoffman, G.E., and Schadt, E.E. (2016). variancePartition: interpreting drivers of variation in complex gene expression studies. BMC bioinformatics 17, 483. 10.1186/s12859-016-1323-z.

39. Satija, R., Farrell, J.A., Gennert, D., Schier, A.F., and Regev, A. (2015). Spatial reconstruction of single-cell gene expression data. Nat Biotechnol 33, 495–502. 10.1038/nbt.3192.

40. La Manno, G., Soldatov, R., Zeisel, A., Braun, E., Hochgerner, H., Petukhov, V., Lidschreiber, K., Kastriti, M.E., Lonnerberg, P., Furlan, A., et al. (2018). RNA velocity of single cells. Nature 560, 494–498. 10.1038/s41586-018-0414-6.

41. Bergen, V., Lange, M., Peidli, S., Wolf, F.A., and Theis, F.J. (2020). Generalizing RNA velocity to transient cell states through dynamical modeling. Nat Biotechnol 38, 1408–1414. 10.1038/s41587-020-0591-3.

42. Braun, E., Danan-Gotthold, M., Borm, L.E., Lee, K.W., Vinsland, E., Lonnerberg, P., Hu, L., Li, X., He, X., Andrusivova, Z., et al. (2023). Comprehensive cell atlas of the first-trimester developing human brain. Science 382, eadf1226. 10.1126/science.adf1226.

43. Fleck, J.S., Sanchis-Calleja, F., He, Z., Santel, M., Boyle, M.J., Camp, J.G., and Treutlein, B. (2021). Resolving organoid brain region identities by mapping single-cell genomic data to reference atlases. Cell stem cell 28, 1148–1159 e1148. 10.1016/j.stem.2021.02.015.

44. Shi, Y., Wang, M., Mi, D., Lu, T., Wang, B., Dong, H., Zhong, S., Chen, Y., Sun, L., Zhou, X., et al. (2021). Mouse and human share conserved transcriptional programs for interneuron development. Science 374, eabj6641. 10.1126/science.abj6641.

45. Conn, P.J., Sanders-Bush, E., Hoffman, B.J., and Hartig, P.R. (1986). A unique serotonin receptor in choroid plexus is linked to phosphatidylinositol turnover. Proc Natl Acad Sci U S A 83, 4086–4088. 10.1073/pnas.83.11.4086.

46. Achim, K., Peltopuro, P., Lahti, L., Tsai, H.H., Zachariah, A., Astrand, M., Salminen, M., Rowitch, D., and Partanen, J. (2013). The role of Tal2 and Tal1 in the differentiation of midbrain GABAergic neuron precursors. Biol Open 2, 990–997. 10.1242/bio.20135041.

47. Goridis, C., and Rohrer, H. (2002). Specification of catecholaminergic and serotonergic neurons. Nat Rev Neurosci 3, 531–541. 10.1038/nrn871.

48. Haldipur, P., Aldinger, K.A., Bernardo, S., Deng, M., Timms, A.E., Overman, L.M., Winter, C., Lisgo, S.N., Razavi, F., Silvestri, E., et al. (2019). Spatiotemporal expansion of primary progenitor zones in the developing human cerebellum. Science 366, 454–460. 10.1126/science.aax7526.

49. Aldinger, K.A., Thomson, Z., Phelps, I.G., Haldipur, P., Deng, M., Timms, A.E., Hirano, M., Santpere, G., Roco, C., Rosenberg, A.B., et al. (2021). Spatial and cell type transcriptional landscape of human cerebellar development. Nat Neurosci 24, 1163–1175. 10.1038/s41593-021-00872-y.

50. Dougherty, K.J., Zagoraiou, L., Satoh, D., Rozani, I., Doobar, S., Arber, S., Jessell, T.M., and Kiehn, O. (2013). Locomotor rhythm generation linked to the output of spinal shox2 excitatory interneurons. Neuron 80, 920–933. 10.1016/j.neuron.2013.08.015.

51. Wang, Y.W., Hu, N., and Li, X.H. (2022). Genetic and Epigenetic Regulation of Brain Organoids. Front Cell Dev Biol 10, 948818. 10.3389/fcell.2022.948818.

52. Lim, Y., Cho, I.T., Shi, X., Grinspan, J.B., Cho, G., and Golden, J.A. (2019). Arx Expression Suppresses Ventralization of the Developing Dorsal Forebrain. Scientific reports 9, 226. 10.1038/s41598-018-36194-6.

53. Kikkawa, T., and Osumi, N. (2021). Multiple Functions of the Dmrt Genes in the Development of the Central Nervous System. Front Neurosci 15, 789583. 10.3389/fnins.2021.789583.

54. Prakash, N., Puelles, E., Freude, K., Trumbach, D., Omodei, D., Di Salvio, M., Sussel, L., Ericson, J., Sander, M., Simeone, A., and Wurst, W. (2009). Nkx6-1 controls the identity and fate of red nucleus and oculomotor neurons in the mouse midbrain. Development 136, 2545–2555. 10.1242/dev.031781.

55. Osterberg, N., Wiehle, M., Oehlke, O., Heidrich, S., Xu, C., Fan, C.M., Krieglstein, K., and Roussa, E. (2011). Sim1 is a novel regulator in the differentiation of mouse dorsal raphe serotonergic neurons. PloS one 6, e19239. 10.1371/journal.pone.0019239.

56. Bayly, R.D., Brown, C.Y., and Agarwala, S. (2012). A novel role for FOXA2 and SHH in organizing midbrain signaling centers. Dev Biol 369, 32–42. 10.1016/j.ydbio.2012.06.018.

57. Lagutin, O.V., Zhu, C.C., Kobayashi, D., Topczewski, J., Shimamura, K., Puelles, L., Russell, H.R., McKinnon, P.J., Solnica-Krezel, L., and Oliver, G. (2003). Six3 repression of Wnt signaling in the anterior neuroectoderm is essential for vertebrate forebrain development. Genes Dev 17, 368–379. 10.1101/gad.1059403.

58. Song, X., Chen, H., Shang, Z., Du, H., Li, Z., Wen, Y., Liu, G., Qi, D., You, Y., Yang, Z., et al. (2021). Homeobox Gene Six3 is Required for the Differentiation of D2-Type Medium Spiny Neurons. Neurosci Bull 37, 985–998. 10.1007/s12264-021-00698-5.

59. Aibar, S., Gonzalez-Blas, C.B., Moerman, T., Huynh-Thu, V.A., Imrichova, H., Hulselmans, G., Rambow, F., Marine, J.C., Geurts, P., Aerts, J., et al. (2017). SCENIC: single-cell regulatory network inference and clustering. Nat Methods 14, 1083–1086. 10.1038/nmeth.4463.

60. Zainolabidin, N., Kamath, S.P., Thanawalla, A.R., and Chen, A.I. (2017). Distinct Activities of Tfap2A and Tfap2B in the Specification of GABAergic Interneurons in the Developing Cerebellum. Frontiers in molecular neuroscience 10, 281. 10.3389/fnmol.2017.00281.

61. Zhang, Y., Rozsa, M., Liang, Y., Bushey, D., Wei, Z., Zheng, J., Reep, D., Broussard, G.J., Tsang, A., Tsegaye, G., et al. (2023). Fast and sensitive GCaMP calcium indicators for imaging neural populations. Nature 615, 884–891. 10.1038/s41586-023-05828-9.

62. Kilb, W. (2021). When Are Depolarizing GABAergic Responses Excitatory? Frontiers in molecular neuroscience 14, 747835. 10.3389/fnmol.2021.747835.

63. Koh, I., and Hagiwara, M. (2023). Gradient to sectioning CUBE workflow for the generation and imaging of organoids with localized differentiation. Commun Biol 6, 299. 10.1038/s42003-023-04694-5.

64. Danesin, C., and Houart, C. (2012). A Fox stops the Wnt: implications for forebrain development and diseases. Curr Opin Genet Dev 22, 323–330. 10.1016/j.gde.2012.05.001.

65. Brodski, C., Blaess, S., Partanen, J., and Prakash, N. (2019). Crosstalk of Intercellular Signaling Pathways in the Generation of Midbrain Dopaminergic Neurons In Vivo and from Stem Cells. J Dev Biol 7. 10.3390/jdb7010003.

66. Chizhikov, V.V., Lindgren, A.G., Currle, D.S., Rose, M.F., Monuki, E.S., and Millen, K.J. (2006). The roof plate regulates cerebellar cell-type specification and proliferation. Development 133, 2793–2804. 10.1242/dev.02441.

67. Lowenstein, E.D., Rusanova, A., Stelzer, J., Hernaiz-Llorens, M., Schroer, A.E., Epifanova, E., Bladt, F., Isik, E.G., Buchert, S., Jia, S., et al. (2021). Olig3 regulates early cerebellar development. eLife 10. 10.7554/eLife.64684.

68. Yeung, J., Ha, T.J., Swanson, D.J., and Goldowitz, D. (2016). A Novel and Multivalent Role of Pax6 in Cerebellar Development. J Neurosci 36, 9057–9069. 10.1523/JNEUROSCI.4385-15.2016.

69. Nakamura, H., Katahira, T., Matsunaga, E., and Sato, T. (2005). Isthmus organizer for midbrain and hindbrain development. Brain Res Brain Res Rev 49, 120–126. 10.1016/j.brainresrev.2004.10.005.

70. Johansen, N., Somasundaram, S., Travaglini, K.J., Yanny, A.M., Shumyatcher, M., Casper, T., Cobbs, C., Dee, N., Ellenbogen, R., Ferreira, M., et al. (2023). Interindividual variation in human cortical cell type abundance and expression. Science 382, eadf2359. 10.1126/science.adf2359.

71. Micali, N., Kim, S.K., Diaz-Bustamante, M., Stein-O’Brien, G., Seo, S., Shin, J.H., Rash, B.G., Ma, S., Wang, Y., Olivares, N.A., et al. (2020). Variation of Human Neural Stem Cells Generating Organizer States In Vitro before Committing to Cortical Excitatory or Inhibitory Neuronal Fates. Cell reports 31, 107599. 10.1016/j.celrep.2020.107599.

72. Gansel, K.S. (2022). Neural synchrony in cortical networks: mechanisms and implications for neural information processing and coding. Front Integr Neurosci 16, 900715. 10.3389/fnint.2022.900715.

73. Berke, J.D., Okatan, M., Skurski, J., and Eichenbaum, H.B. (2004). Oscillatory entrainment of striatal neurons in freely moving rats. Neuron 43, 883–896.

74. Klaus, A., and Plenz, D. (2016). A Low-Correlation Resting State of the Striatum during Cortical Avalanches and Its Role in Movement Suppression. PLoS Biol 14, e1002582. 10.1371/journal.pbio.1002582.

75. Plenz, D. (2003). When inhibition goes incognito: feedback interaction between spiny projection neurons in striatal function. Trends Neurosci 26, 436–443.

## Supplemental References

1. Braun, E., Danan-Gotthold, M., Borm, L.E., Lee, K.W., Vinsland, E., Lonnerberg, P., Hu, L., Li, X., He, X., Andrusivova, Z., et al. (2023). Comprehensive cell atlas of the first-trimester developing human brain. Science 382, eadf1226. 10.1126/science.adf1226.

2. Fleck, J.S., Sanchis-Calleja, F., He, Z., Santel, M., Boyle, M.J., Camp, J.G., and Treutlein, B. (2021). Resolving organoid brain region identities by mapping single-cell genomic data to reference atlases. Cell stem cell 28, 1148–1159 e1148. 10.1016/j.stem.2021.02.015.

3. Aldinger, K.A., Thomson, Z., Phelps, I.G., Haldipur, P., Deng, M., Timms, A.E., Hirano, M., Santpere, G., Roco, C., Rosenberg, A.B., et al. (2021). Spatial and cell type transcriptional landscape of human cerebellar development. Nat Neurosci 24, 1163–1175. 10.1038/s41593-021-00872-y.

4. Aibar, S., Gonzalez-Blas, C.B., Moerman, T., Huynh-Thu, V.A., Imrichova, H., Hulselmans, G., Rambow, F., Marine, J.C., Geurts, P., Aerts, J., et al. (2017). SCENIC: single-cell regulatory network inference and clustering. Nat Methods 14, 1083–1086. 10.1038/nmeth.4463.

